# CRISPR screens establish regulatory maps of immunosuppressive surface molecules in cancer

**DOI:** 10.64898/2026.03.10.710555

**Authors:** Robert Kalis, Sumit Deswal, Markus Schäfer, Mathias Kalxdorf, Julian Jude, Jesse Lipp, Sarah Rieser, Vivien Vogt, Melanie de Almeida, Michaela Fellner, Stefanie Ruhland, Lisa Frasz, Florian Andersch, Jeroen Krijgsveld, Sebastian Carotta, Johannes Zuber

**Affiliations:** Research Institute of Molecular Pathology (IMP), Vienna BioCenter (VBC), 1030 Vienna, Austria; Vienna BioCenter PhD Program, Doctoral School of the University of Vienna and Medical University of Vienna; Division Proteomics of Stem Cells and Cancer, German Cancer Research Center (DKFZ), Heidelberg, Germany; Boehringer Ingelheim RCV, 1120 Vienna, Austria; Medical Faculty, Heidelberg University, Heidelberg, Germany; Medical University of Vienna, Vienna BioCenter (VBC), 1030 Vienna, Austria

## Abstract

Cancer cells can evade immune surveillance by triggering inhibitory checkpoint responses in tumor-associated T cells through the expression of immunosuppressive surface molecules. While therapeutic blockade of such receptors has emerged as a pillar of cancer therapy, tumor cell-intrinsic mechanisms controlling their expression remain incompletely understood. Fluorescence-activated cell sorting (FACS)-based genetic screens can be used to decipher regulatory pathways, but conventional screening approaches are biased towards regulators that are dispensable for cancer cell proliferation and survival. Here, we used a tetracycline-inducible Cas9 system enabling fully time-controllable CRISPR-based mutagenesis to gain a more comprehensive and comparative survey of regulators controlling the expression of four major immunosuppressive surface molecules, PD-L1 (CD274), CD47, CD276 and HLA-E, as well as CD151, a candidate surface target associated with tumor growth and invasion. As a prominent hit, our screens identify the membrane-trafficking factor DNAJC13 as a regulator of PD-L1 and CD276. Among DNAJC13-controlled surface proteins, we identify other known and proposed immune-checkpoint molecules. Based on this function, suppression of DNAJC13 strongly increases the sensitivity of human cancer cells to T-cell attack *in vitro* and prolongs survival of mice bearing pancreatic tumors. Together, our study establishes regulatory maps of major immune-modulatory surface molecules and identifies DNAJC13 as a potential target for the coordinated inhibition of multiple immunosuppressive signals.

## Introduction

A major immune evasion mechanism of cancer cells is the expression of immune-modulatory surface molecules that trigger inhibitory checkpoint responses in tumor-associated immune cells. Therapeutic antibodies blocking such signals can restore anti-tumor immunity and have revolutionized cancer medicine, as exemplified by the groundbreaking success of PD-1/PD-L1 blocking antibodies in the treatment of advanced melanoma and other cancers^1,2^. PD-L1 belongs to a group of immune-modulatory surface molecules that act as ligands of B7 and CD28 receptors on T-cells. Expression of PD-L1 (CD274, B7-H1) on tumor cells and accessory cells in the tumor microenvironment engages checkpoint responses in tumor-specific T-cells upon binding to the PD-1 (PDCD1) receptor^3,4^. Other immune checkpoints currently considered for therapeutic exploitation include the ‘do not eat me signal’ receptor CD47, a major regulator of innate immune surveillance^5,6^. CD47 is frequently expressed on cancer cells to suppress tumor-associated macrophages via binding to inhibitory transmembrane receptor SIRPα present on myeloid cells. CD276 (B7-H3) is overexpressed in a wide range of human cancers and correlates with poor clinical outcome, yet its exact function and the corresponding receptor remain unknown^7,8^. HLA-E is a non-classical MHC class I molecule that is overexpressed in many cancers and has been shown to inhibit the activity of NK and CD8+ T-cells by binding to NKG2A/KLRC1^9^. Blocking this immune checkpoint has recently been shown to trigger potent antitumor responses^10,11^.

While the groundbreaking success of PD-1/PD-L1 checkpoint blockers has led to a paradigm shift in cancer medicine, clinical progress has – at least to some degree – outpaced our understanding of the underlying biology. A better understanding of mechanisms and pathways controlling the expression of immunosuppressive surface molecules in normal tissues and cancer is crucial to guide the clinical use of different checkpoint blockade therapeutics and may reveal new targets for immune-modulatory therapies. Combining functional-genetic screens with flow cytometry-based readouts and fluorescence-activated cell sorting (FACS) has emerged as a powerful approach for deciphering such regulatory networks. Both genome-scale CRISPR/Cas9- and haploid gene-trap screens have been used to investigate regulators of PD-L1, leading to the identification of CMTM6 as a critical regulator controlling PD-L1 recycling^12,13^. While the convergent identification of CMTM6 using independent cellular models and screening methods highlights its importance in PD-L1 regulation, the overall number of regulators identified in these screens is surprisingly low. One limitation of these and other previous ‘regulator screens’ is that they relied on stable gene knockout and involved prolonged periods of time for cell selection and expansion between mutagenesis and the flow-cytometry based readout. During this time, cells with knockout events that compromise cellular fitness are lost from the cell population, so such screens may miss important regulators.

To enable a more systematic, truly genome-wide analysis of regulatory pathways, we combined a doxycycline (Dox)-inducible Cas9 expression system^14^ and a second-generation sgRNA library^15^ to perform a series of time-controlled FACS-based CRISPR screens. Results of these screens establish regulatory maps of major immunosuppressive surface molecules in cancer that, besides known non-essential factors, uncover previously unknown regulators regardless of their effect on cell proliferation and survival. Among them, we identify DNAJC13 as a trafficking factor that coordinates the surface expression of several immunosuppressive cues.

## Results

### CRISPR/Cas9 screens identify regulators of PD-L1 surface expression

To gain insight into cancer-cell intrinsic mechanisms controlling PD-L1 expression, we sought to perform FACS-based CRISPR screens in a cell line that expresses PD-L1 in the absence of interferon γ (IFN-γ), which is known to induce PD-L1 in most cell types^16^. By re-analyzing RNA sequencing data of 675 human cancer cell lines^17^ we found exceptionally high *PD-L1* expression levels in RKO cells (Extended data Fig. 1a), a mismatch-repair deficient colorectal cancer cell line harboring microsatellite instability (MSI), which is associated with responsiveness to PD-1/PD-L1 checkpoint blockade in patients^18^. RKO cells have acquired a super-enhancer around the PD-L1 transcriptional start site (Extended data Fig. 1b and c) that likely drives constitutive PD-L1 expression, but otherwise harbor a near-diploid genome without major copy number aberrations. These features and their robust growth in culture made RKO cells ideally suited for exploring cancer cell-intrinsic mechanisms driving PD-L1 expression.

To gain full temporal control over CRISPR mutagenesis, we introduced a two-vector Dox-inducible Cas9 expression system (iCas9) in RKO cells (Fig. 1a), isolated single-cell clones via FACS, and tested Cas9 expression and function in expanded clones by transducing lentiviral vectors co-expressing a fluorescent marker and an sgRNA targeting PLK1, a generally essential kinase, or PD-L1. In the clone selected for subsequent studies, prolonged culture in the absence of Dox did not result in a depletion of sgPLK1-expressing cell (Fig. 1b) and no reduction of PD-L1 in sgCD274-expressing cells (Fig. 1c), demonstrating that the activity of Cas9 is strictly Dox-dependent. Dox treatment led to a rapid, near-complete depletion of sgPLK1-expressing cells (Fig. 1b, Extended data Fig. 1d) and a complete loss of PD-L1 surface expression in sgCD274-expressing cells (Fig. 1c), indicating that Dox-inducible CRISPR mutagenesis is highly effective using the iCas9 system.

**Figure 1.**
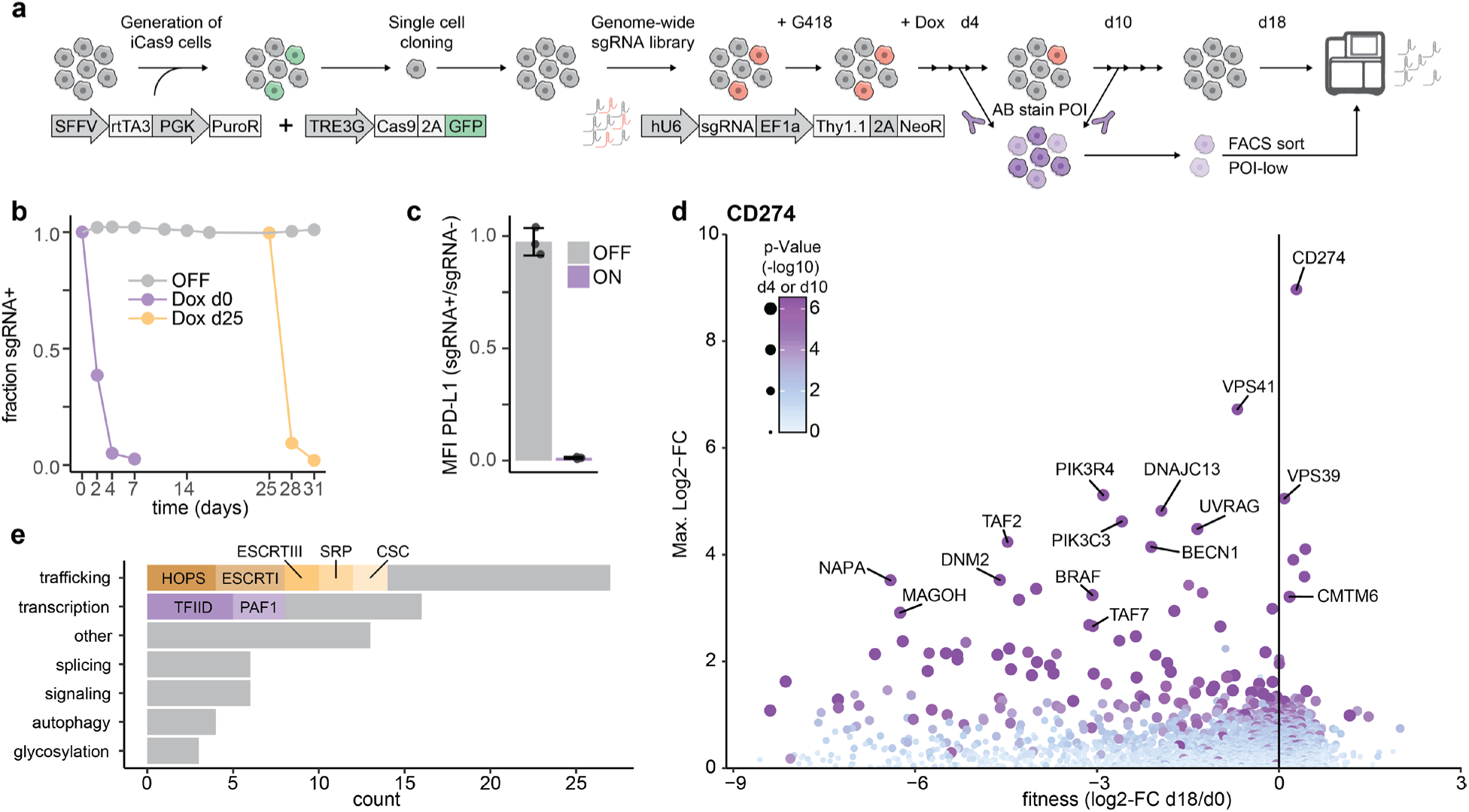
CRISPR screen for regulators of PD-L1 (CD274). **a**, Schematic of Tet-Cas9 cell line engineering and FACS-based CRISPR screens for PD-L1 regulators. **b**, Competitive proliferation assay of Tet-Cas9 RKO cells transduced with an sgRNA targeting PLK1 in the presence or absence of Dox. The percentage of sgRNA+ cells was monitored by flow cytometry for 31 days after Cas9 induction. **c**, Flow cytometry-based quantification of PD-L1 expression (median fluorescence intensity, MFI) following inducible editing of PD-L1. **d**, FACS-based CRISPR screens for positive regulators of PD-L1 surface expression. For each gene, the maximum effect of two sorting timepoints (average sgRNA enrichment in sorted PD-L1 low cells compared to the starting point, d0) is plotted against the depletion (average sgRNA fold-change after 18 days of proliferation compared to d0). Dot size and color indicate the p-value (-Log10) of the respective plotted timepoint. The negative y-axis is not shown. **e**, Annotation of the top 75 enriching genes (by Log2-fold change, p-Value < 0.01) based on gene function and association with indicated protein complexes.

In addition to PD-L1 itself, we investigated several established regulators of PD-L1 using single sgRNA-assays in iCas9-RKO cells, including components of the IFN-γ pathway (IRF1, STAT1, CDK5)^16,19^, CMTM6^12,13^, and mutant BRAF(V600E)^20^, which acts as a driving oncogene in RKO cells^21^. Inducible knockout of IRF1, STAT1 and CDK5 did not result in any reduction of PD-L1 surface levels, indicating that the high constitutive expression of PD-L1 is completely independent of the IFN-γ pathway in RKO cells (Extended data Fig. 1e). Knockout of BRAF and CMTM6 led to a reduction in PD-L1 surface levels, in line with their previously described roles in PD-L1 regulation.

For genome-wide screening, we transduced iCas9-RKO cells with the Vienna genome-wide CRISPR library^15^ at high representation of approximately 1000 cells/sgRNA. Tight control over Cas9 allowed us to drug select, expand, and cryopreserve library-transduced iCas9-RKO cells in large quantities prior to CRISPR mutagenesis, and thereby create a renewable resource for genome-wide CRISPR screens. To systematically identify PD-L1 regulators, we reasoned that differences in mRNA and/or protein stability, as well as drop-out effects due to essential gene functions, could be accounted for by sorting cells displaying reduced PD-L1 levels at two different time points following Cas9 induction. To identify regulators independent of essential gene functions, on day 4 following Cas9 induction we sorted the 1% of cells showing the lowest PD-L1 levels, and performed a second sort using a more stringent gating strategy (lowest 0.1%) on day 10 to be able to capture more slowly turned-over regulators (Extended data Fig. 2a). In addition, we maintained Dox-treated iCas9-RKO cells in culture for 18 days to perform a conventional drop-out screen and thereby assess essential gene functions and the overall quality of our screen. Comparing the representation of sgRNAs in Dox-treated cells following 18 days of culture (T18) to cells prior to Dox induction (T0) revealed a strong depletion of previously annotated ‘core essential’ genes^22^ and other dependencies in RKO cells (Extended data Fig. 2b). Notably, with an average depletion of >30-fold, the drop-out of core essential genes exceeded effect sizes observed in most previous drop-out screens (Extended data Fig. 2c), indicating that CRISPR mutagenesis is highly efficient in the iCas9-RKO model.

In cells sorted for low PD-L1 expression, all sgRNAs targeting CD274 were the strongest enriched on both day 4 and day 10 (Extended data Fig. 2d, Extended data Table 1) and, consequently, CD274 was the top-scoring hit in an integrated analysis of both time points (Fig. 1d). Other prominent hits included CMTM6 and BRAF, both showing strong effects already after 4 days of Cas9 induction. In line with the dependency of RKO cells on BRAF^V600E^, sgRNAs targeting BRAF strongly depleted over time, indicating that our screening approach captured PD-L1 regulators that are required for cell proliferation and survival. In contrast to previous CRISPR-based and haploid gene-trap screens, which revealed CMTM6 as the only prominent hit besides CD274 and IFN-γ pathway components^12,13^, our time-resolved screening strategy identified 154 genes showing significant sgRNA enrichment (p<0.01, LFC>1) in cells sorted for low PD-L1 expression (Fig. 1d, Extended data Table 1). Among these, 56 genes showed a marked sgRNA depletion over time in our drop-out screen (p<0.01, LFC<-2), indicating that many PD-L1 regulators are also required for cellular fitness. These even included broadly essential genes such as NAPA and CDC73, which were typically enriched in PD-L1 low cells at the early time point and then strongly depleted. This explains why many hits were missed in previous screens involving prolonged culture prior to FACS and demonstrates that our screening approach identifies regulators independent of essential cellular functions.

The list of identified PD-L1 regulators encompasses factors involved in diverse cellular processes, from transcriptional gene control, including several components of the general transcription factor TFIID, and mRNA splicing to intracellular protein trafficking and turnover (Fig. 1e). Prominent hits included several genes implicated in endocytic trafficking such as all four components of the ESCRTI complex (TSG101, VPS37, VPS28, UBAP1)^23^. We further identify major components of the HOPS complex (VPS41, VPS39, VPS33A and VPS18)^24^ and its interactor RAB7A^25^ to be required for PD-L1 expression. Other notable hits with a described protein trafficking function include PIK3C3/PIK3R4, UVRAG/BECN1, and DNAJC13. While several of these factors have been shown to physically and functionally interact in endosome trafficking and autophago-lysosome maturation^26,27^, results of our screens suggest that they may cooperate in an endocytic recycling pathway that enhances PD-L1 surface expression.

To validate our screening results and quantify the impact of regulators acting at different levels on PD-L1 surface expression, we performed a series of single sgRNA assays. In addition to iCas9-RKO cells, we included the human pancreatic adenocarcinoma cell line MIA-PaCa2 as an independent cellular model, for which we also derived a Dox-inducible Cas9 clone (iCas9-MP). Since MIA-PaCa2 cells express very low levels of PD-L1 at baseline (Extended data Fig. 1a), validation studies were performed upon treatment with IFN-γ, which strongly induces PD-L1 expression in iCas9-MP cells and provided an additional regulatory context. In total, we selected 24 prominent screen hits, for each of which transduced 2-3 scoring sgRNAs individually into iCas9-RKO and -MP cells, and quantified changes in PD-L1 surface expression in sgRNA-expressing cells 4 and 10 days following Cas9 induction (Fig 2a-b, Extended data Fig. 3a-f). All selected hits validated to regulate PD-L1 surface expression in iCas9-RKO cells with a high correlation of maximum effect sizes at the two time points (Pearson r = 0.77; Fig. 2c). Most of these hits also validated to be required for PD-L1 expression in iCas9-MP cells upon IFN-γ induction with a high-correlation of effect sizes (Pearson r = 0.64; Fig. 2d, Extended data Fig. 3g). Notable differences between the two cell lines include BRAF, which does not act as a driving oncogene and had only minor effects on PD-L1 expression in iCas9-MP cells, as well as PIK3C3/PIK3R4, UVRAG and BECN1, which showed delayed and attenuated effects in comparison to iCas9-RKO cells (Fig. 2b). Taken together, these validation experiments confirmed that our screening approach provides a comprehensive survey of previously unknown, broadly relevant PD-L1 regulators.

**Figure 2.**
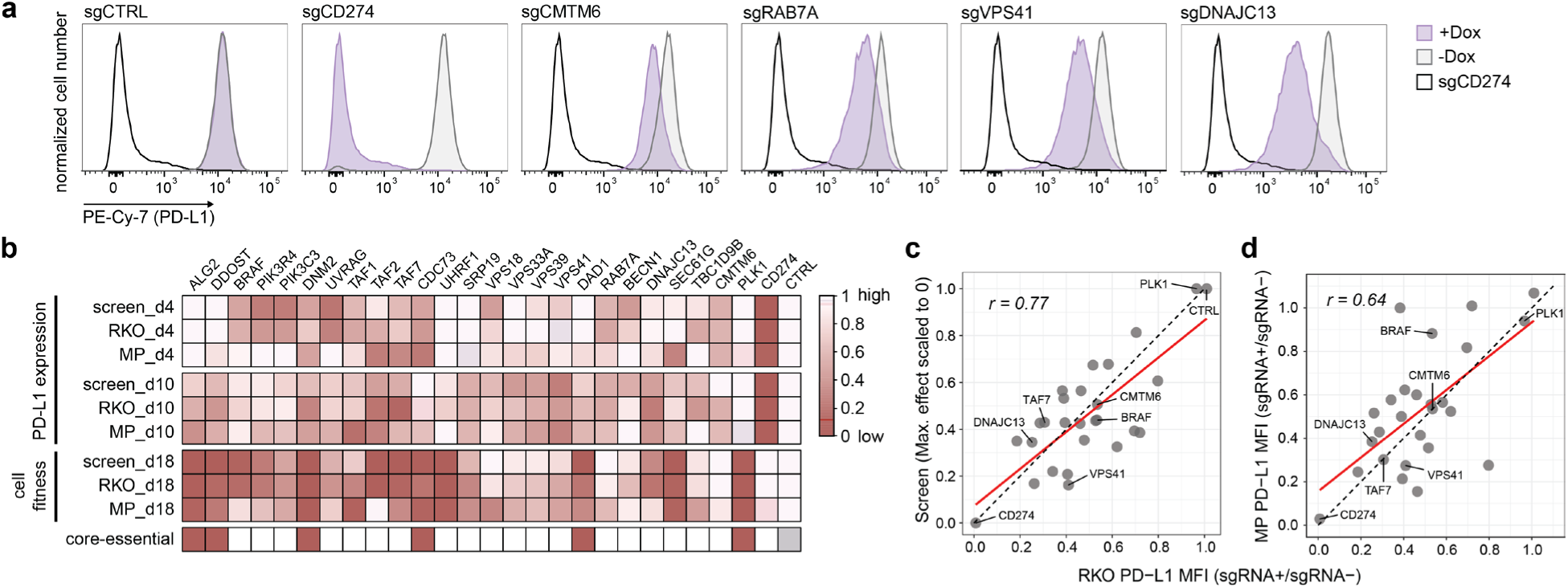
Extended validation of 24 regulators of PD-L1 surface expression. **a**, Flow cytometry quantification of PD-L1 expression after inducible editing of selected screen hits and controls. Surface PD-L1 was quantified by immunostaining and flow cytometry 10 days after Cas9 induction. Data is representative of three independent experiments with three different sgRNAs per gene. **b**, Heatmap of effects on PD-L1 surface expression and cell proliferation in single sgRNA assays in two independent cell lines (RKO, MP = MIA PaCa-2 pre-treated with 10 ng/ml IFN-y) and the original screen. For each gene, cells were transduced with individual sgRNA constructs with an infection efficacy of 30-60%. The effect of 2 or 3 different guide RNAs per gene was averaged and scaled based on suitable controls: PD-L1 expression levels (upper panels) in the screen were scaled between no sgRNA enrichment (Log2FC ≤ 0, set to 1 = control PD-L1 expression) and the effect after PD-L1 knockout (average of all 5 sgRNAs, set to 0 = low PD-L1 expression). In single sgRNA assays, PD-L1 expression was quantified based on the ratio of median fluorescence intensities (MFI) in sgRNA-expressing and non-expressing cells within each sample (MFI sgRNA+ / MFI sgRNA-) on day 4 (d4) and day 10 (d10) following Cas9 induction. This comparison assumes cell-autonomous effects but better controls for cell density. Cellular fitness (lower panels) in the screen was scaled between no sgRNA depletion (Log2-FC ≥ 0, set to 1 = high fitness) and the average effect of ‘core-essential’ genes (Log2-FC = −4, set to 0 = low fitness). In single sgRNA assays in RKO and MP cells, cellular fitness indicates the relative ratio of sgRNA expressing cells after 18 days of culture, compared to the initial fraction prior to Cas9 induction. Core-essential: 0 indicates core-essential genes. **C - d**, Scatter plot comparing averaged sgRNA effects on PD-L1 in single guide validation studies in RKO cells to the screen (c) and to MP cells (d). The maximum effect on either day 4 or day 10 is shown. r, Pearson correlation; red line indicates linear fitting model calculated in R; dashed black line indicates x = y.

### FACS-based CRISPR screens establish regulatory maps of other immune-modulatory surface proteins and CD151

After establishing a scalable workflow for FACS-based screens, we extended this approach to investigate regulators of other immunomodulatory surface molecules including CD47, CD276 and HLA-E to enable comparative analyses and explore potential co-regulators. As a fifth surface factor, we included CD151, a structurally and functionally unrelated transmembrane protein implicated in invasion and metastasis^28,29^, which in pilot experiments was rapidly lost upon CRISPR-mediated knockout (Extended data Fig. 4a), suggesting that CD151 is dynamically regulated and rapidly turned over. For the purpose of comparative screens, CD151 therefore provides a suitable control to distinguish between more selective or more general regulators in protein expression and trafficking.

Expression of all four proteins in RKO cells was confirmed in RNA-sequencing^14^ and flow cytometry (Extended data Fig. 4b-c). For genome-wide screens, we took advantage of library transduced iCas9-RKO cells that had been cryo-preserved following G418 selection prior to Cas9 induction at high representation (>1000 cells/sgRNA). Deep sequencing of sgRNA cassettes after thawing showed that prolonged culture in the absence of Dox and a freeze-thaw cycle did not lead to major representation shifts or a dropout of sgRNAs targeting essential genes (Extended data Fig. 4d). This confirms the tightness of our iCas9 system at library scale, enabling the use of cryopreserved library-transduced cells as renewable resource for consecutive screens. Genome-wide FACS-based screens for all four additional factors were performed analogous to the PD-L1 screen, with the exception that we sorted 1% of cells with the lowest expression at both time points (Extended data Fig. 4e). In all four screens, the surface protein under investigation was identified as a top-scoring gene among numerous regulators, many of which are required for cellular fitness based on our drop-out screen (Fig. 3a-d). The number of identified regulators varied between screens. The least were identified in the HLA-E screen, most likely due to the relatively low dynamic range of fluorescent signal after HLA-E antibody staining (Extended data Fig. 4c and e). Nevertheless, several specific regulators of HLA-E could be identified, including Beta-2 Microglobulin (B2M), which is required for surface expression of all HLA class I heavy chains^30^. The highest number of hits was obtained in the CD151 screen, which - at least in part - could be attributed to the rapid turnover of this surface molecule as this facilitates the detection of regulators with essential cellular functions. In case of more stable surface proteins, knockout of essential regulators can lead to a dropout or more complex phenotypes before effects on the expression of the surface protein become detectable. Hence, we would not consider all regulators identified only in the CD151 screen to be CD151-specific. Conversely, selective regulators identified in other screens that do not reduce CD151 levels are unlikely to be generally required for protein expression or trafficking.

**Figure 3.**
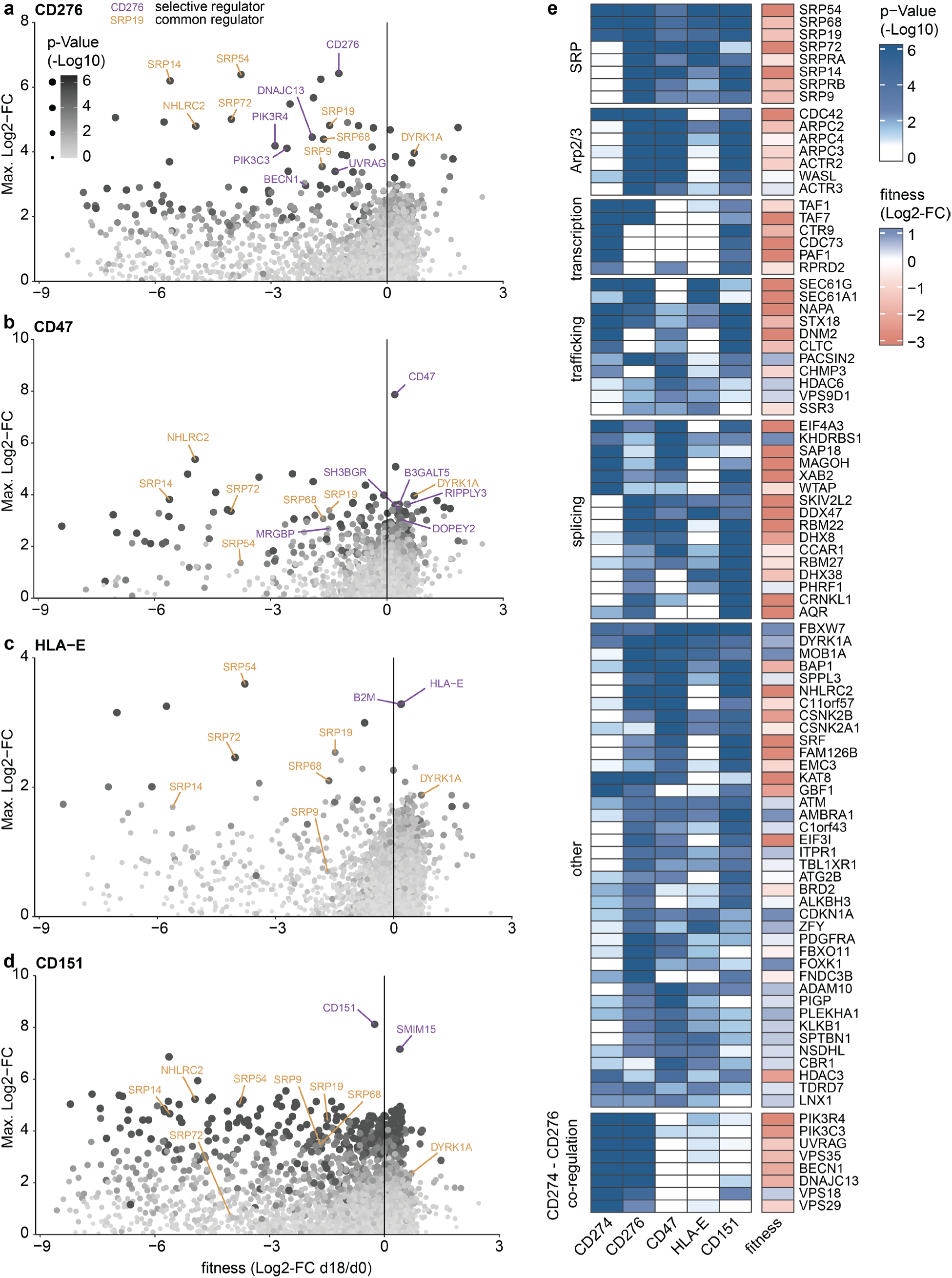
CRISPR screens for regulators of CD276, CD47, HLA-E, and CD151. **a – d**, FACS-based CRISPR screens for positive regulators of the surface molecules CD276 (a), CD47 (b), HLA-E (c), and CD151 (d). For each gene, the maximum effect of the two sorting timepoints (average sgRNA enrichment in sorted POI-low cells compared to starting point, d0) is plotted against depletion effects (average ratio of sgRNA abundance at the d18 endpoint compared to d0). Dot size and color indicate p-value (based on MAGeCK; -Log10) of the respective plotted timepoint. Selective regulators, defined as enriching significantly in only one of the four screens, are highlighted in purple; common regulators, defined as enriching in at least three screens, are highlighted in orange. Negative y-axis is not shown. **e**, Heatmap of all selected genes enriching significantly (p-Value < 0.01) in at least two screens. Color indicates p-Value (-Log10). ‘fitness’ denotes depletion effects shown in (a – d).

Taking these potential limitations into account, we compared the results of the five FACS-based screens and identified 169 regulators that scored in two or more screens with high confidence (p<0.01, LFC>2; Fig. 3e, Extended data Fig. 5a-d). Among such common regulators, we identified all six protein components and both receptor subunits of the signal recognition particle (i.e. SRP9, SRP14, SRP19, SRP54, SRP68, SRP72, SRPA and SRPB), a highly conserved ribonucleoprotein complex broadly required for targeting nascent proteins to their proper membrane localization^31^. In addition, we identified all five nonredundant subunits of the ARP2/3 complex (ACTR2, ACTR3, APRC2, ARPC3, ARPC4), a central regulator of the actin cytoskeleton, as well as CDC42 and WASL, which directly regulate the ARP2/3 complex in processes such as endocytosis and Golgi trafficking^32^. Other common regulators include components of core transcriptional machinery and the spliceosome (Extended data Fig. 5b-c), which showed the most pronounced effects in the CD151 screen while most likely being required for the expression of all investigated surface molecules.

Taken together, our screens represent a single, comprehensive CRISPR-based dataset focused on surface protein expression. A particularly interesting group of regulators had strong effects on PD-L1 and CD276 expression, while being dispensable for other surface molecules including CD151, indicating that they are not generally required for surface protein expression. These include the PIK3C3/R4-UVRAG-BECN1 complex and DNAJC13, which was among the top hits in both the PD-L1 and the CD276 screen (Fig. 1d, Fig. 3a). This prompted us to further investigate DNAJC13, which might be exploitable for inhibiting multiple immunosuppressive signals on cancer cells without globally interfering with the surface proteome.

### DNAJC13 is a context-specific cancer dependency

DNAJC13 (DnaJ Heat Shock Protein Family Member C13; also known as RME-8) acts as co-chaperone of a partner heat-shock protein (HSC70)^33^. Mechanistically, DNAJC13 is known to localize to the endosome membrane, where it interacts with the retromer and WASH complexes^34^. DNAJC13 has been implicated as a regulator of endosomal subdomain organization, protein trafficking and sorting, but its specific functions in mammalian cells remain incompletely understood^35,36^. In RKO cells, our screen and validation experiments indicate that loss of DNAJC13 confers modest effects on cellular fitness (Extended data Fig. 3e). To assess whether this reflects a generally essential function, we queried DNAJC13 alongside two similarly scoring PD-L1 regulators, TAF7 and CMTM6 (Fig. 2b), across 612 CRISPR screens^15,37–44^ that recalled ‘core essential genes’ with a high dynamic range (LFC<-2, on average). TAF7 drops out in most screens, in line with its role as a component of the general transcription factor TFIID, whereas such effects were only observed in very few cell lines for CMTM6 (Fig. 4a). DNAJC13 turns out to be required in some but dispensable in most cell lines. Indeed, inducible knockout of DNAJC13 had a minor impact on cell proliferation or survival in Mia-Paca2, A375 melanoma, or 639V urothelial carcinoma cells, while we observed a modest sgRNA depletion in RKO and MDAMB231 breast cancer cells over 10 days of culture (Fig. 4b). By contrast, PD-L1 expression levels were strongly reduced in all tested cell lines (Fig. 4c), indicating that the context-specific dependency on DNAJC13 is independent of its role in PD-L1 regulation.

**Figure 4.**
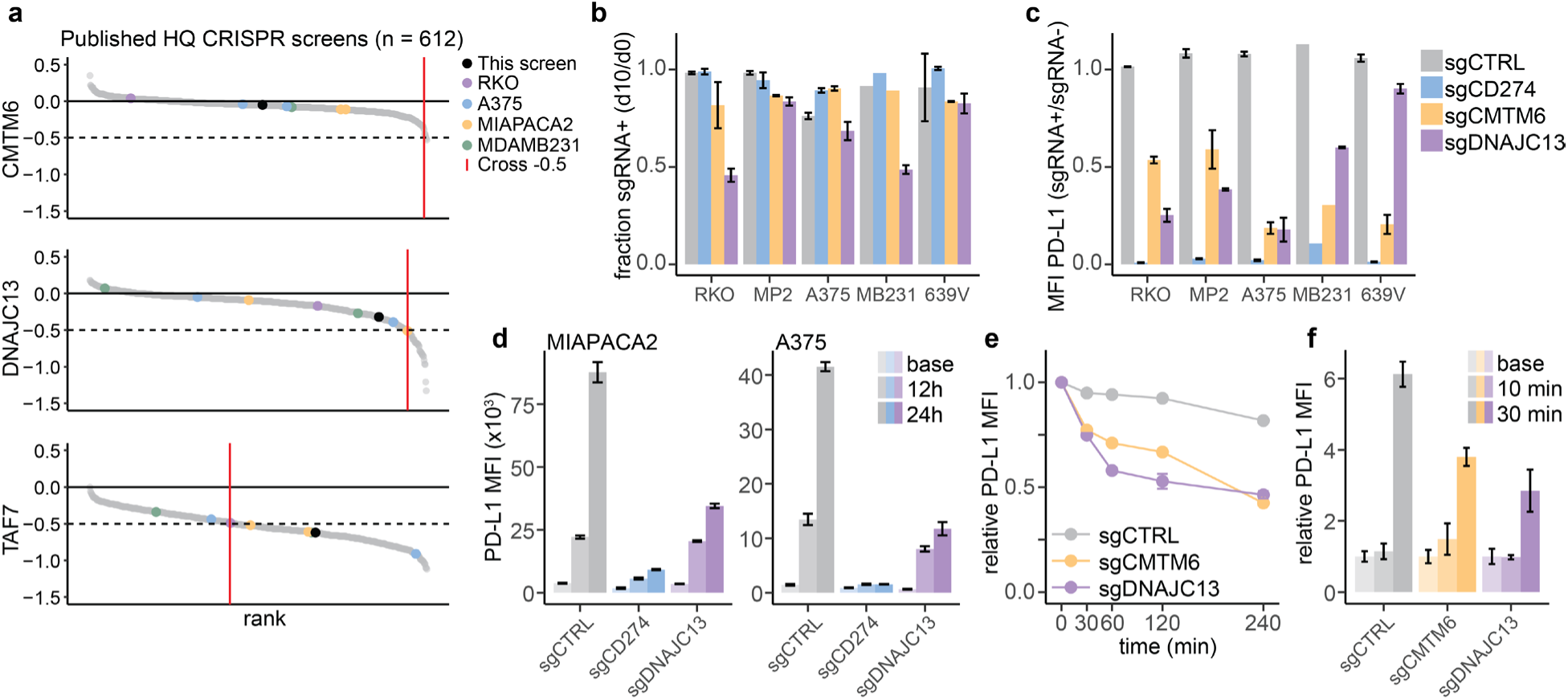
DNAJC13 is a context-specific dependency and post-translational PD-L1 regulator. **a**, Depletion effects of indicated genes in 612 published genome-wide CRISPR screens, scaled to −1 for core-essential genes and 0 for non-essential genes. Cell lines used for validation experiments are highlighted. **b**, Fitness effect upon KO of DNAJC13 in cancer cell lines from diverse tissue contexts. MP2 = Mia-PaCa2, MB231 = MDA-MB-231. Cells transduced with indicated sgRNAs were cultured for 10 days following Cas9 induction, and the fraction of sgRNA+ cells was quantified relative to the initial population prior to Cas9 induction. **c**, MFI (Median Fluorescence Intensity) of antibody-stained PD-L1 was compared in sgRNA+ and sgRNA-cells of the same sample (MFI sgRNA+ / MFI sgRNA-) 10 days after Cas9 induction. MIA PaCa2 and A375 cells were pre-treated with IFN-y. The assay was performed in the same experiment as shown in (b). Experiments in (b - c) were conducted in independent duplicates (except MDA-MB231). Two different DNAJC13 guides were used in each duplicate, error bars indicate SD. **d**, MIAPACA2 cells (left) or A375 cells (right) harboring perturbations of Control, CD274 or DNAJC13 were treated with 10ng/ul IFNy, and PD-L1 MFI was assessed over time by flow cytometry. **e**, A375 cells were stained with unconjugated IgG-anti-PD-L1 antibody, thoroughly washed and then placed at 37 degrees. PD-L1 MFI was assessed by anti-IgG staining at indicated timepoints. **f**, A375 cells were stained with unconjugated IgG-anti-PD-L1 antibody, thoroughly washed and then placed at 37 degrees for 60 minutes to allow internalization. Cells were then washed with a low pH stripping buffer to remove surface bound antibody. Cells were again placed at 37 degrees to allow re-expression of internalized (and stripping-protected) antibody-bound PD-L1. At indicated timepoint, re-expression was assessed by anti-IgG staining and flow-cytometry. For d-e, n = 3 independent replicates.

### DNAJC13 is a post-translational regulator of several immune-modulatory surface proteins

DNAJC13 has been shown to regulate EGFR surface trafficking^45^ and, more recently, has been implicated as a positive modulator of autophagy^36^. Thus, it seemed most likely that DNAJC13 controls PD-L1 at the post-translational level during membrane trafficking or post-endocytic recycling, as has been described for CMTM6^12,13^. In line with a post-translational function, loss of CMTM6 or DNAJC13 did not alter PD-L1 mRNA levels, in contrast to the knockout of TAF7 or BRAF, and did not lead to any major global changes in mRNA expression (Extended data Fig. 6a-b). To probe such a role, we determined how the deletion of DNAJC13 affects the kinetics of PD-L1 induction upon IFN-y treatment. Deletions of PD-L1 itself led to strongly reduced PD-L1 surface levels at 12h and 24h timepoints following Cas9 induction, whereas such effects only unfolded at 24h upon loss of DNAJC13 (Fig. 4d). This and the DNAJC13 protein localization to the endosomal compartment (Extended data Fig. 6c), indicates that DNAJC13 is not required for initial expression of PD-L1, and prompted us to investigate its role in membrane-PD-L1 recycling. Both RKO and A375 cells harboring a DNAJC13 deletion lost surface PD-L1 faster than control cells, to a similar extent as upon deletion of the known PD-L1 stabilizer CMTM6 (Fig. 4e, Ext. Fig 6d). DNAJC13 deficient cells were also compromised in their ability to re-express internalized IgG-stained PD-L1 (Fig. 4f, Extended data Fig. 6e). Together, this suggests that DNAJC13 functions in membrane protein recycling.

As the knockout of DNAJC13 or CMTM6 only led to a partial loss of PD-L1 surface expression, we investigated whether top screen hits, particularly those implicated in endocytic trafficking, would synergize with DNAJC13 in PD-L1 regulation. Knockout of a second regulator enhanced effects on PD-L1 expression in both DNAJC13 and CMTM6 KO backgrounds (Extended data Fig. 6f). However, even then, cells retained PD-L1 expression, suggesting that endocytic trafficking pathways boost - but are not essential for - PD-L1 surface expression.

To study the consequences of DNAJC13 loss in an unbiased fashion, we performed mass spectrometry-based proteomics in DNAJC13 or control KO RKO cells. Total proteomics showed the depletion of DNAJC13 as well as PD-L1 and PD-L2 in DNAJC13 edited cells (Extended data Fig. 7a). Interestingly, the set of 130 significantly downregulated proteins is highly enriched for surface molecules (Extended data Fig. 7b). Prompted by these results, we sought to increase sensitivity by performing surface protein-enrichment prior to mass spectrometry in DNAJC13, CMTM6 and control KO cells. As RKO cells exhibit a modest growth disadvantage upon DNAJC13 loss, we included the melanoma cell line A375 to independently validate our results. This approach robustly identified around 750 cell surface proteins per sample. Consistent with screening and validation results, the levels of PD-L1 and CD276 in DNAJC13 deficient A375 cells were significantly depleted (Fig. 5a-b). In comparisons between the effects of CMTM6 and DNAJC13 KO in A375 cells, we find a surprisingly high correlation of effect sizes (r = 0.56; Fig. 5c), potentially indicating redundant mechanisms. Furthermore, several major effects upon DNAJC13 loss, including depletion of PD-L1, were consistent between RKO and A375 cells (r = 0.36; Fig. 5d, Extended data Fig. 7c-d). We observed an upregulation of several proteins involved in metabolic processes, such as glycolysis or ribosephosphate-synthesis, as well as an upregulation of proteins associated with intracellular and endosomal transport (Extended data Fig. 7e-g). Gene Set Enrichment Analysis confirmed consistent effects of DNAJC13 KO, both in RKO and A375, on cell-adhesion and immune-related proteins (Fig. 5e-f; Extended data Fig. 7h). Besides PD-L1 and CD276, we find a significant depletion of NT5E/CD73, a complex associated with the production of immunosuppressive adenosine metabolites^46^, as well as PVR and PVRL2, two ligands of T-cell immunoreceptor with Ig and ITIM domains (TIGIT), a known target for checkpoint inhibition^47^. Taken together, we find that DNAJC13 regulates a subset of surface proteins that are enriched for immune-related molecules but does not affect many other surface molecules or MHC class I (Extended data Fig. 7i-k).

**Figure 5.**
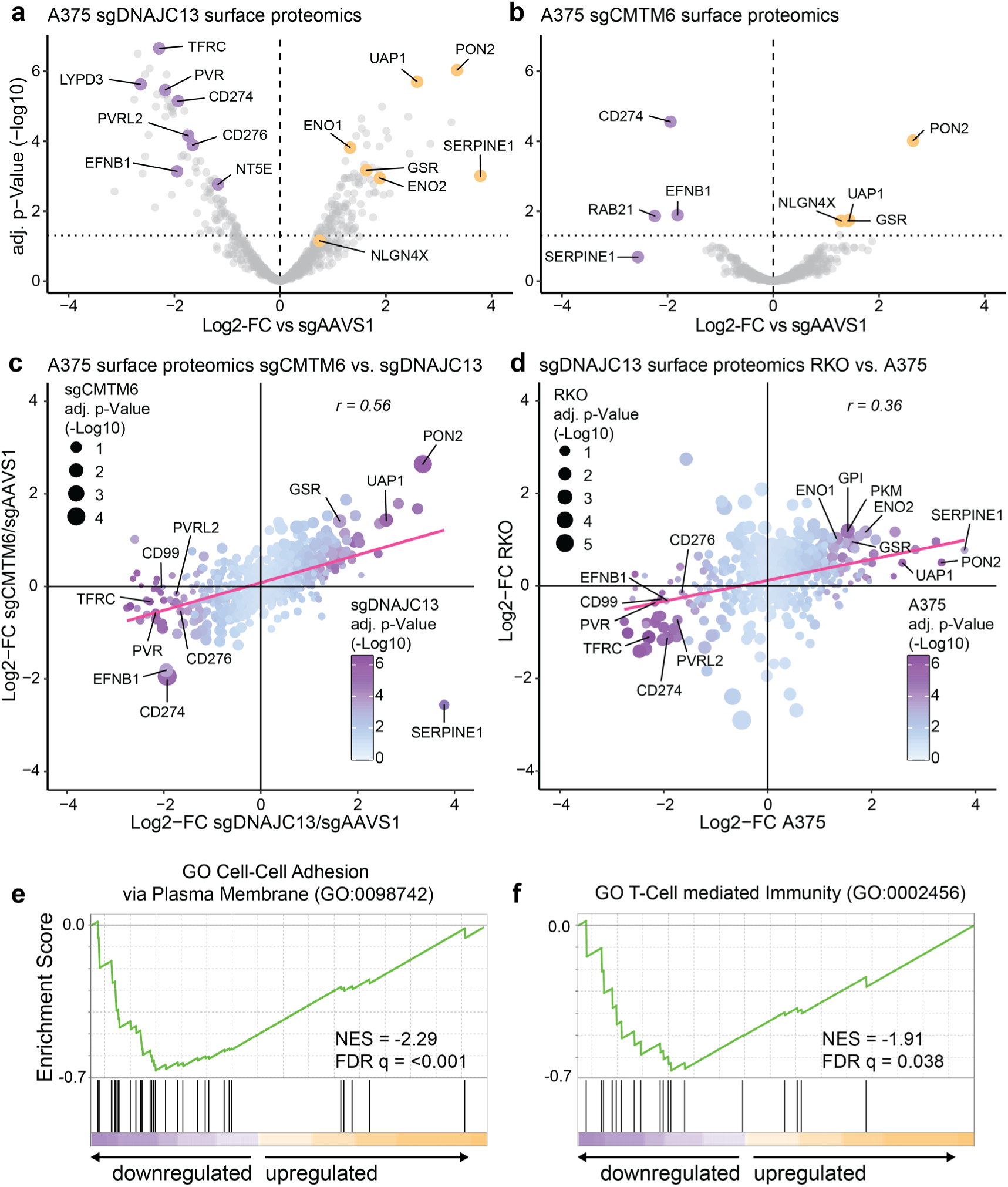
Systematic analysis of surface proteins controlled by DNAJC13 and CMTM6. **a - b**, Surface proteomics in A375 cells expressing sgDNAJC13 (a) or sgCMTM6 (b), in comparison to sgAAVS1 neutral control. Cells were cultured for 8 days following Cas9 induction and treated with 10 ng/ml IFN-y for 48h prior to sampling. **c**, Comparison of effects following CRISPR-mediated knockout of CMTM6 versus DNAJC13 in A375 cells (based on data in a-b). Dot size indicates adj. p-values in the sgCMTM6 sample; color codes indicate adj. p-values in the DNAJC13 sample. **d**, Comparison of Log2FC of sgDNAJC13 expressing A375 and RKO cells (see also Extended data Fig. 7c). Dot size indicates adj. p-values in the RKO sample; color code indicates adj. p-values in the A375 sample. Red line displays linear fitted model; r represents Pearson correlation (c – d). **e - f**, Gene set enrichment analysis (GSEA) of sgDNAJC13-induced effects in A375 cells. Log2-fold changes (vs. sgAAVS1) of all detected surface proteins were processed in a weighted rank-based GSEA to compute enriched gene sets. NES = normalized enrichment score; p-values in panels a-d were calculated using LIMMA with Benjamini-Hochburg correction.

### Loss of DNAJC13 renders cancer cells sensitive to T-cell mediated killing

To test whether the coordinated deregulation of these immune-modulatory surface molecules translates into increased susceptibility to T-cell attack, we devised a co-culture assay between HLA-A*0201-expressing iCas9-RKO cells and primary human T-cells expressing a transgenic T-cell receptor (CMV1-TCR) that recognizes the HLA-A*0201-bound cytomegalovirus peptide NLV^48^ (Extended data Fig. 8a). Pulsing of HLA-A*0201 transgenic iCas9-RKO with the NLV peptide led to a marked reduction of cancer cell numbers upon co-culture with CMV1-TCR transgenic T-cells (Extended data Fig. 8b). While loss of PD-L1 did not augment these effects, knockout of DNAJC13 markedly increased the susceptibility of iCas9-RKO cells to T-cell-mediated killing (Fig. 6a).

**Figure 6.**
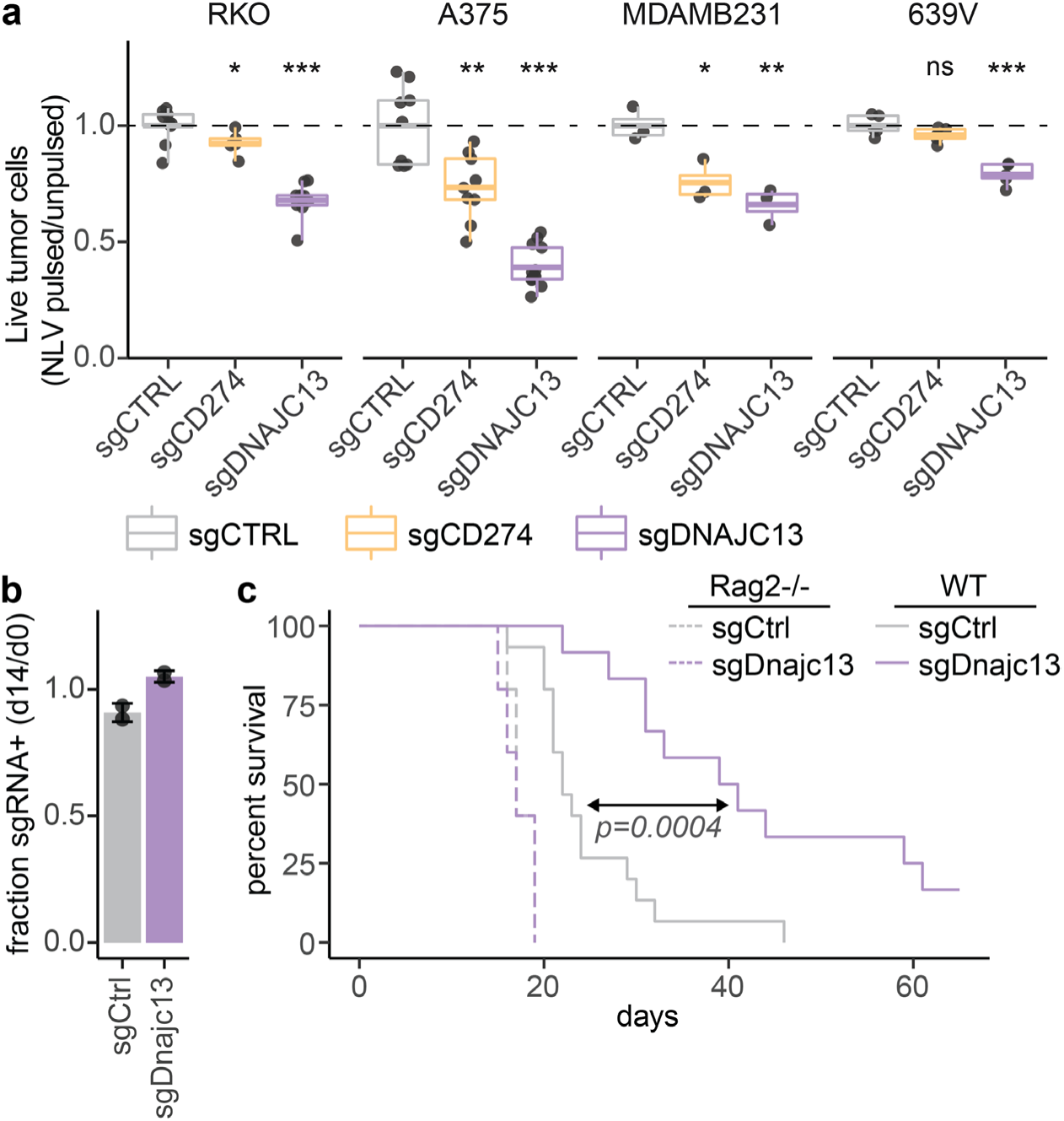
Suppression of DNAJC13 enhances CD8^+^ T cell-mediated tumor cell killing. **a**, Comparison of live tumor cell counts in NLV-pulsed samples and unpulsed conditions in RKO, A375, MDA-MB-231, and 639V cells. A375 and 639V were pre-treated with 10ng/ml IFN-y 48h prior to addition of T-cells. Effect sizes were normalized to the respective sgAAVS1 sample. Points show individual values from replicates (n = 3–5) with a line indicating the mean and error bars from Min. to Max. P-values were calculated using unpaired t-tests with Welch correction, with * = p < 0.05, ** = p < 0.005, *** = p < 0.0005. **b**, in vivo competition assay of sgCtrl or sgDnajc13 cells in orthotopic EPP2 pancreas tumors in B6 Rag2-/- mice. **c**, survival of B6 Rag2-/- or B6-WT mice bearing orthotopic EPP2 pancreas tumors harboring deletions of Control or Dnajc13 (n=5 for Rag2-/-, n=15 (control) or n=12 (Dnajc13) for WT). WT graphs show an integration of 3 independent experiments. p-value was calculated using unpaired t-test with Welch correction. Cas9 was induced 5 days prior to transplantation in both (b) and (c).

To exclude that these effects are specific to RKO cells or in any way biased by the fitness defect observed upon DNAJC13 KO in this cellular context, we repeated these experiments in additional cancer cell lines that endogenously express HLA-A*0201 and are insensitive to loss of DNAJC13. In all cases, loss of DNAJC13 strongly increased the susceptibility of cancer cells to T-cell mediated killing (Fig. 6a, Ext. Fig 8c-d). Strikingly, the impact of DNAJC13 KO on T-cell mediated killing was more pronounced than upon knockout of PD-L1, suggesting that these effects are not only a consequence of PD-L1 downregulation but also due to a reduction of other immune-suppressive surface molecules. To test this, we engineered RKO and A375 cells to harbor individual or combinatorial deletions of 5 major DNAJC13-regulated immunosuppressive surface molecules (PD-L1, CD276, PVR, NECTIN2 and NT5E) and directly compared their susceptibility to T-cell mediated killing. While individual perturbations had no or moderate impact, the combinatorial deletion of these 5 molecules closely mirrored effects observed upon DNAJC13 deletion, suggesting that a concerted downregulation of multiple immune checkpoints mediates the effects of DNAJC13 suppression (Extended data Fig. 8e-f).

To study consequences of DNAJC13 loss in a more physiological setting, we utilized a murine model of pancreatic cancer^49^. EPP2 tumor cells can be manipulated in cell culture and readily re-transplanted into the pancreas of recipient mice, where they can also be tracked via Luciferase bioluminescence imaging. Dnajc13 abrogation did not display any proliferation defects in iCas9-EPP2 cells in culture or effects on tumor growth upon transplantation in immunodeficient Rag2-/-mice (Fig. 6b, Extended data Fig. 8g). In striking contrast, immunocompetent mice bearing Dnajc13 deleted tumors showed a significantly delayed tumor progression and improved survival (Fig. 6c, Extended data Fig. 8h), implicating Dnajc13 as a potential target for cancer immunotherapies.

## Discussion

Previous FACS-based screens investigating the regulation of PD-L1 and other surface molecules have relied on stable loss-of-function perturbations and prolonged culture periods, which inevitably leads to negative selection of genes whose knockout compromises the cellular fitness prior to the readout. Consequently, results are biased towards non-essential genes and against genes with additional, potentially unrelated regulatory functions required for cell proliferation and/or survival. We overcame this limitation by employing a Tet-inducible Cas9 clonal cell line that allowed us to detect effects on surface protein levels of dozens of genes that would likely be missed in conventional CRISPR screening approaches. Another major technical advantage of our tightly inducible Cas9 expression is the ability to expand and cryo-preserve sgRNA library-transduced cells prior to genome editing without major changes in library representation, thereby providing a renewable resource for subsequent screens. We used this approach to comparatively survey factors controlling the surface expression of four immunomodulatory surface molecules expressed in cancer, as well CD151. Results of our comparative screens substantially expand the list of PD-L1 and CD47 regulators identified in previous screens^12,13,50,51^ and provide insights into common and specific pathways in surface protein expression.

As one of the most prominent regulators of several immunosuppressive surface factors we identify DNAJC13 as a mediator of PD-L1 and CD276 surface expression. Prior studies have implicated DNAJC13 as a regulator of endosomal trafficking, while certain mutations have been linked to a familial form of Parkinson’s disease^52,53^. Depletion of DNAJC13 has been shown to reduce surface levels of EGFR and transferrin receptor in mammalian cells^35,45^. DNAJC13 has further been described as a positive modulator of autophagy^36^ and recently been detected in a CRISPR screen as a positive regulator of PD-L1^50^. Mechanistically, DNAJC13 associates with early endosomes and is known to interact with the WASH and Retromer complexes^45,54^. In line, we find VPS35 and VPS29, components of the retromer subcomplex CSC (cargo-selective complex) among co-regulators of PD-L1 and CD276, whereas none of the WASH complex members score, possibly due to genetic redundancies. Given our genetic data and previous reports, we hypothesize that loss of DNAJC13 leads to endosomal retention and/or misrouting to degradative pathways. The strong effect on PD-L1 expression and the consistency of phenotypes observed upon knockout of DNAJC13 and CMTM6 also raises the possibility that DNAJC13 acts in the same pathway and functionally cooperates with CMTM6 to protect PD-L1 from lysosomal degradation^12^.

In comparison to CMTM6, loss of DNAJC13 leads to more complex changes in the cell surface proteome that include a coordinated down-regulation of several known and proposed immune-checkpoint triggers such as PD-L1, CD276, PVR/PVRL2 and NT5E. While being well-tolerated in most cancer cell lines under standard culture conditions, knockout of DNAJC13 strongly enhances their sensitivity towards T-cell mediated killing. The scale of these effects exceeds and thus cannot be explained by the loss of PD-L1 alone, and is indeed approximated by simultaneous deletion of 5 different DNAJC13-regulated immunosuppressive surface molecules. This suggests that the immune-sensitized state upon suppression of DNAJC13 is a consequence of a coordinated downregulation of multiple immunomodulatory surface molecules. Its potential as immunotherapy target is underscored by significantly increased survival of mice bearing Dnajc13 deleted pancreatic tumors. Although therapeutic suppression of DNAJC13 may have more severe consequences than in cultured cell lines, essential organismal functions do not preclude its suitability as a target, as demonstrated by many cancer therapeutics in clinical use. Moreover, cancer-cell selective drug delivery approaches such as antibody-drug conjugates could help to mitigate unintended organismal drug activities. Based on our findings, we propose that transient suppression of DNAJC13 may open opportunities to sensitize cancer cells for a wide spectrum of established and emerging cancer immunotherapies.

## Acknowledgments

We are grateful to all members of the Zuber lab for experimental advice and helpful discussions. We thank K. Aumayr, G. Schmauss, M. Weninger and the IMP/IMBA BioOptics team for cell sorting; The IMP/IMBA protein biochemistry core facility for performing quantitative proteomics; A. Sommer and the VBCF-NGS team (www.vbcf.ac.at) for deep sequencing services; H. Scheuch, R. Heinen and the IMP/IMBA molecular biology service for continuous support.

## Author contributions

R.K., S.D. and J.Z. conceived the study and planned this project; R.K. and S.D. designed and conducted experiments with help from M.S., S.R., V.V., St.R., L.F. and M.dA.; R.K., S.D., M.S. and J.Z. analyzed and interpreted original data; M.K. performed mass spectrometry and MS data analysis; J.J. and M.F. cloned the sgRNA library; R.K. performed data analysis with support from J.L. and F.A.; J.J. and M.S. established critical reagents and methodology; J.K. and S.C. provided critical input on experimental designs and data analyses; R.K. and J.Z. wrote the manuscript and all authors reviewed the manuscript.

