## Supplementary Materials for "CRISPR screens establish regulatory maps of immunosuppressive surface molecules in cancer"

R. Kalis et al.

### Extended data Figures

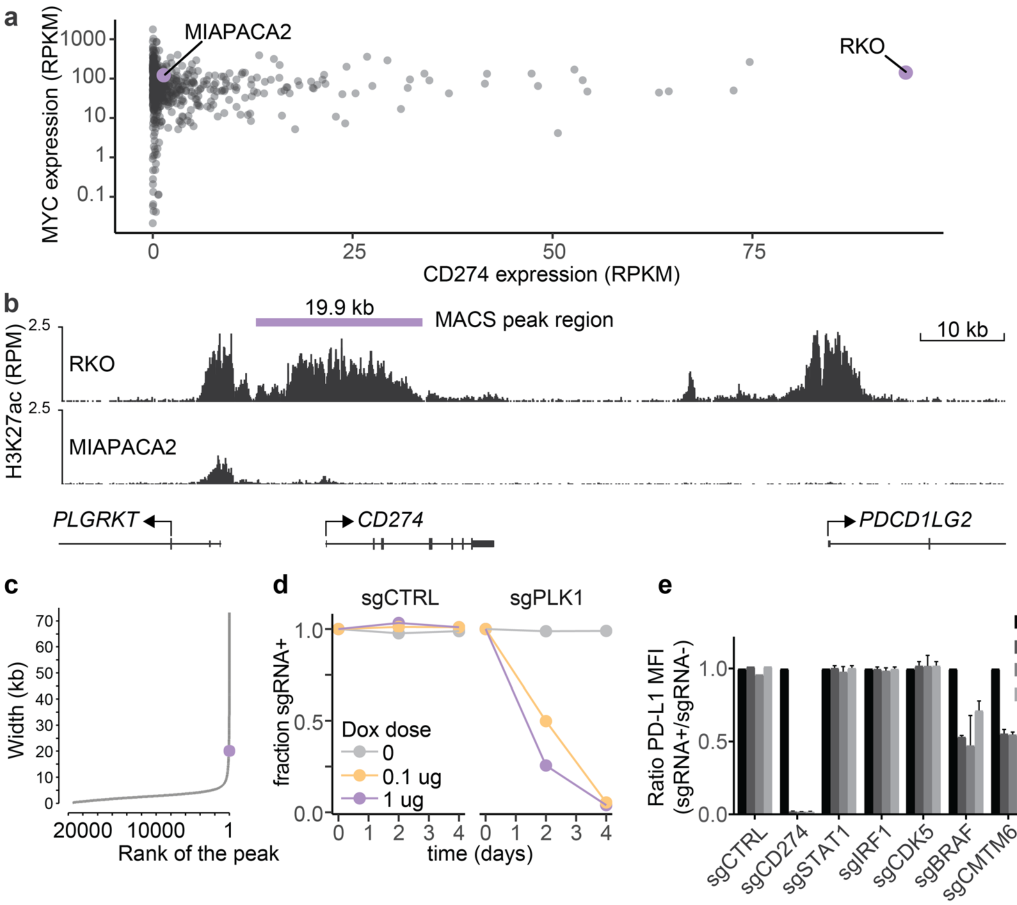


**Extended data Fig. 1 | PD-L1 is highly and constitutively expressed in RKO. a**, mRNA expression of PD-L1 (CD274) plotted against MYC mRNA expression (RPKM). Data is taken from Klijn et al.[17] **b,** RPM-normalized H3K27ac ChIP-seq data at the CD274 locus in RKO and MIAPACA2 cells. **c,** The rank (by peak width) of 21402 H3K27ac peaks identified in RKO cells is plotted against the peak width (kb), with the 19.9 kb CD274 MACS peak region shown in b highlighted in red at rank 81. **d**, Competitive proliferation assay of Tet-Cas9 RKO cells transduced with an sgRNA targeting PLK1 or CTRL, in the presence of indicated doses of Dox (ug/ml). The percentage of sgRNA+ cells was monitored by flow cytometry. **e**, MFI (Median Fluorescence Intensity) of antibody-stained PD-L1 was compared in sgRNA+ and sgRNA- cells of the same sample (MFI sgRNA+ / MFI sgRNA-) at several timepoints after inducible editing.

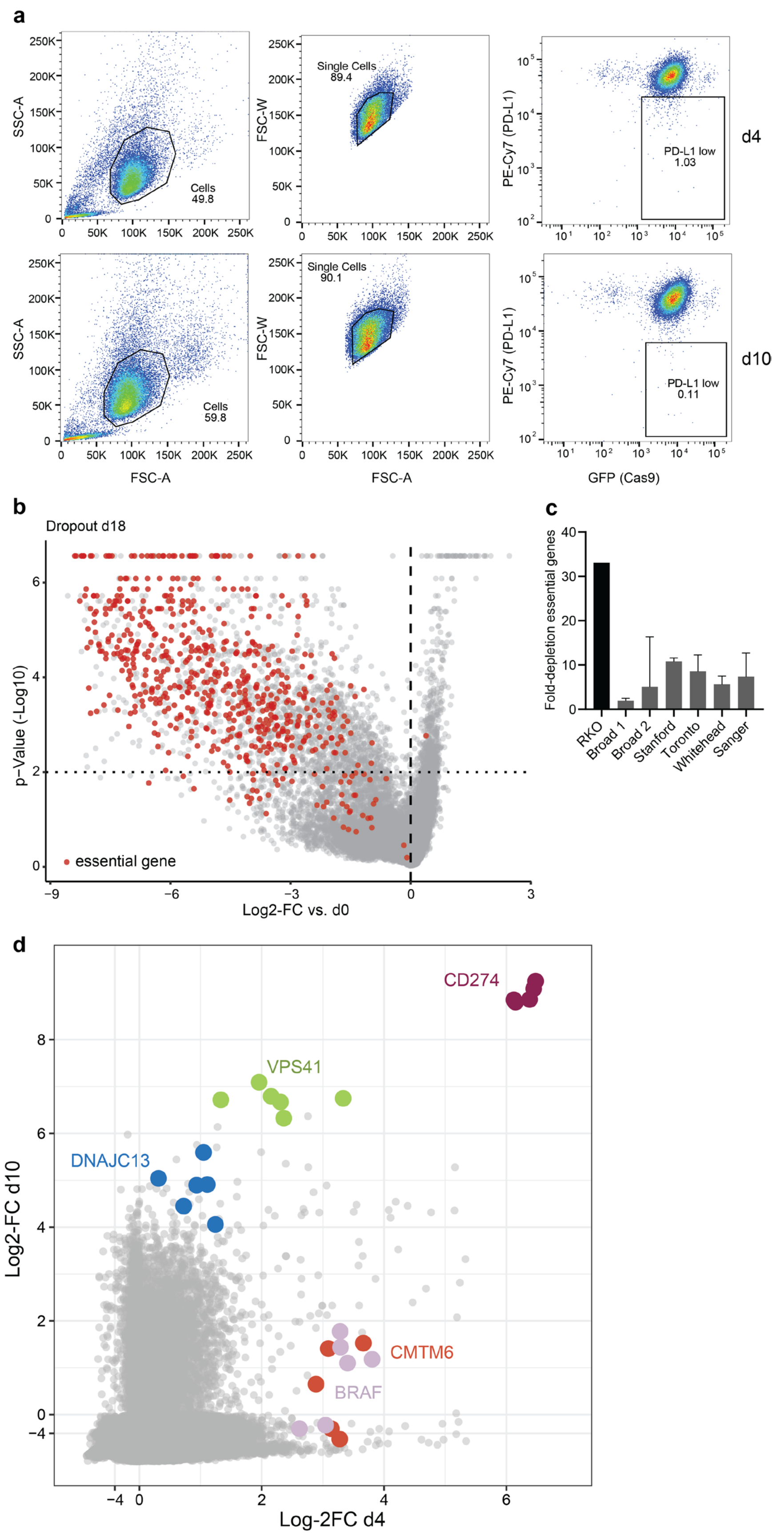


**Extended data Fig. 2 | Dropout results and gating strategy of the time-resolved CRISPR screen. a**, Gating strategy for FACS sorting on days 4 and 10 after inducible editing. **b**, Volcano plot of depletion values of all genes. Core-essential genes highlighted in red. **c**, Re-analysis of published CRISPR screens using MAGeCK[57] and binning according to publications. The average fold-depletion of each bin is plotted with error bars indicating standard deviation. **d**, Guide level results of the PD-L1 screen.

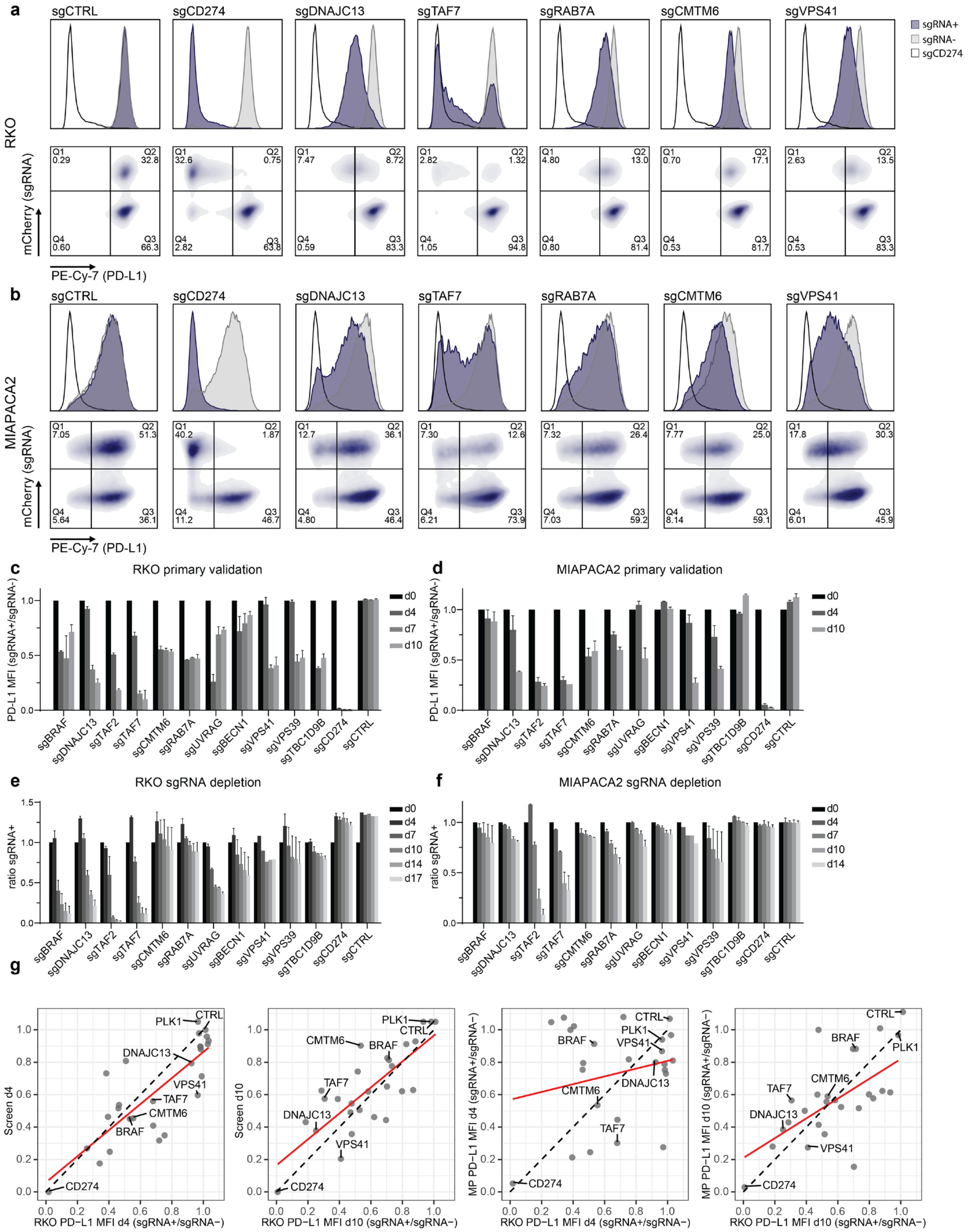

**Extended data Fig. 3 | Validation of PD-L1 screen hits in RKO and MiaPaCa2. a – b** Representative Histograms of RKO cells (a) and MIAPACA2 (b) transduced with sgRNAs targeting selected screen hits. PD-L1 levels were assessed 10 days after Cas9 induction by flow cytometry. MIAPACA2 were pre-treated with IFNy. Representative plots from Fig. 2 (a – b) are shown. **c - d**,MFI of antibody stained PD-L1 was compared in sgRNA+ and sgRNA- cells of the same sample (MFI sgRNA+ / MFI sgRNA-) 10 days after inducible editing in RKO (c) and MIAPACA2 (d). MIAPACA2 cells were pre-treated with IFN-y. **e - f**, Fitness effect upon KO of selected genes in RKO (c) and MIAPACA2 (d). The assay was performed in the same samples as in (b - c). **g**, Scatter plot comparing averaged sgRNA effects on PD-L1 in single guide validation studies in RKO cells to the screen (left) and to MP cells (right), data from Fig 2c-d, split by timepoints.

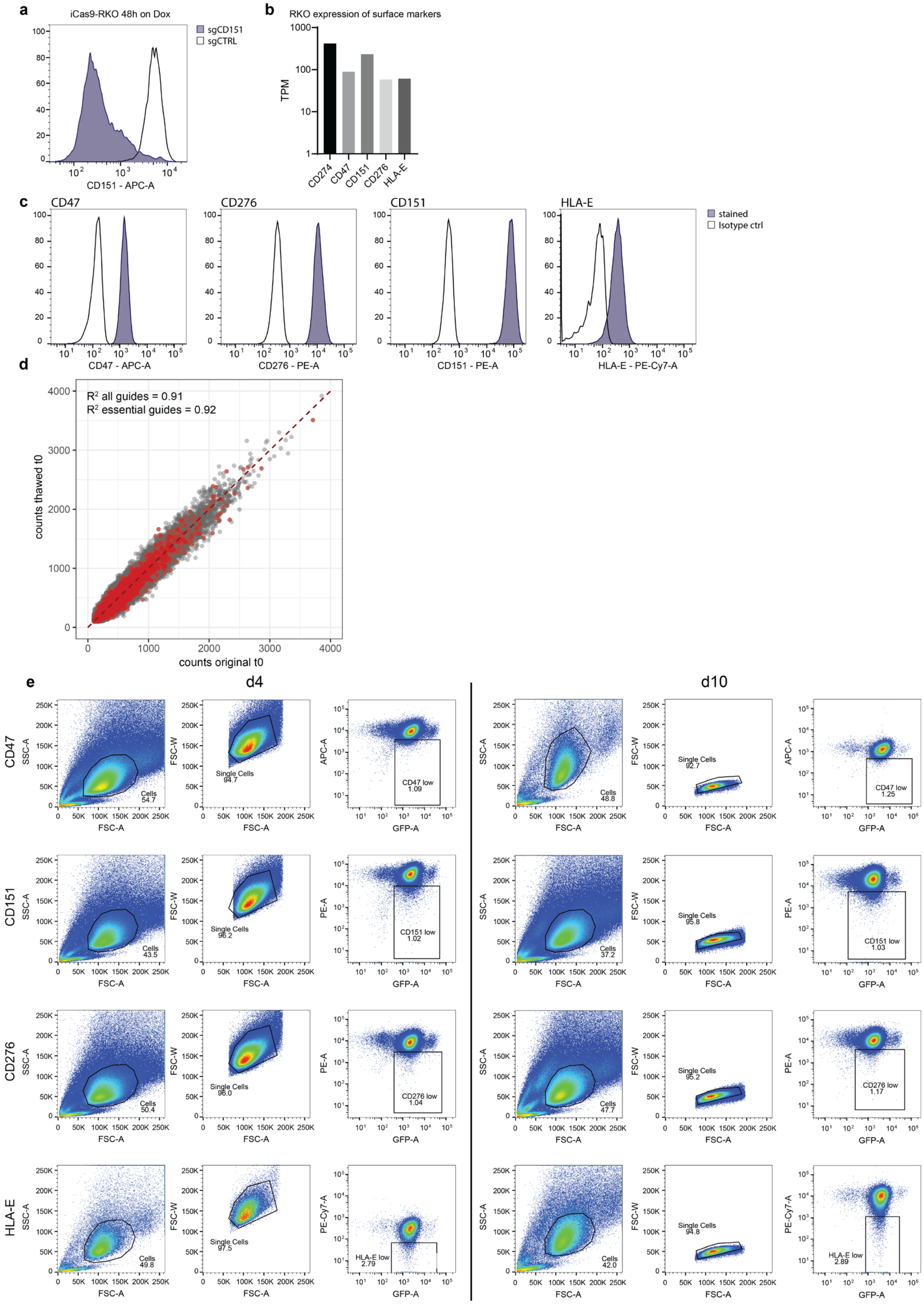

**Extended data Fig. 4 | Expression of screened surface markers and gating strategy for screen sorts. a**, Histograms of CD151 expression in RKO-iCas9 48 hours after inducible editing. **b**, Expression of selected surface molecules in RKO cells. **c**, Histograms of Tet-Cas9 clonal RKO cells stained for expression of selected surface molecules. **d**, Scatter plot of normalized guide counts (filtered for guides >50 counts) comparing the original t0 and t0 after freeze-thaw and prolonged culture **e**, Gating strategy for FACS-sorts on days 4 and 10.

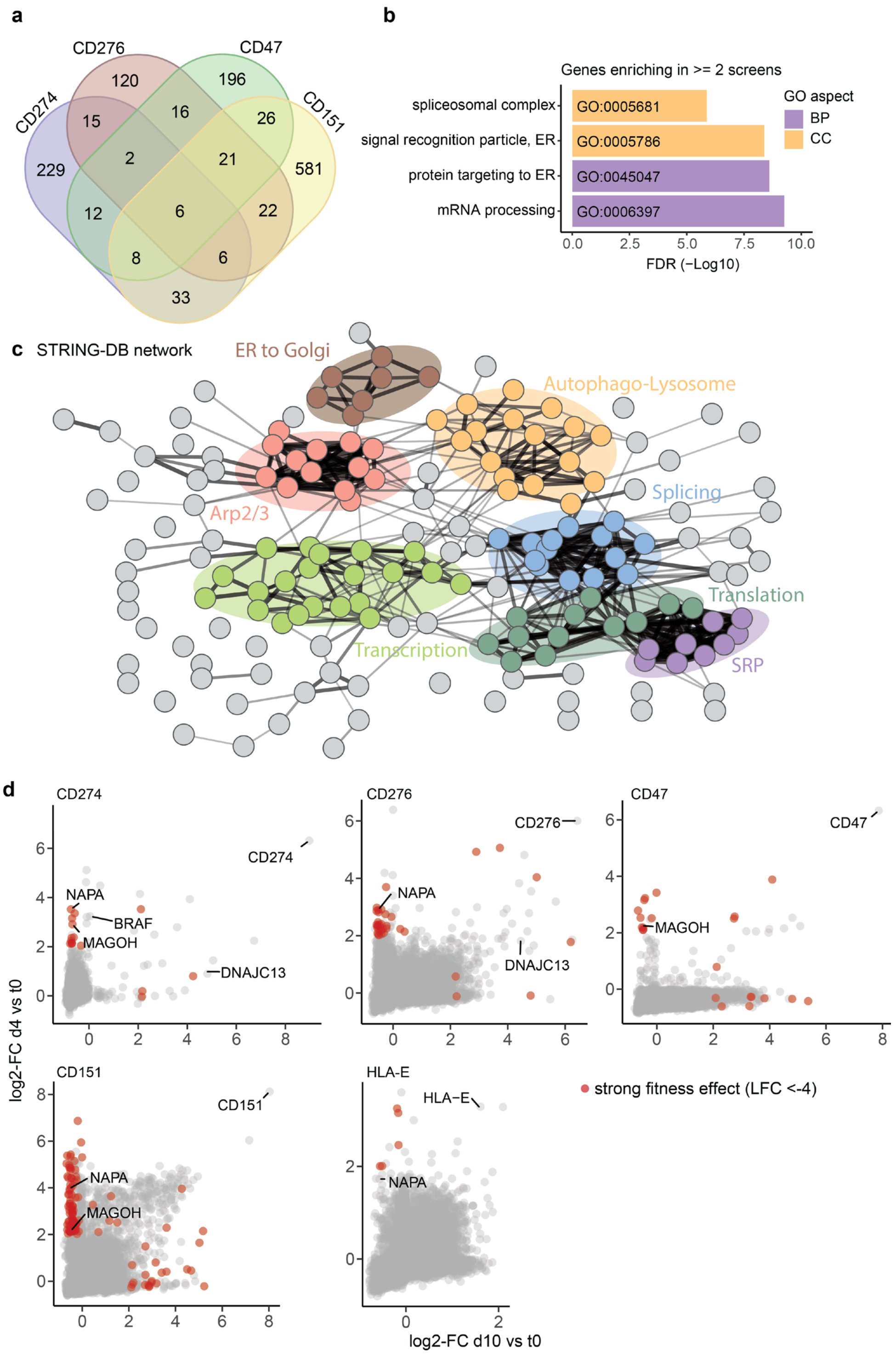


**Extended data Fig. 5 | Meta-analysis of identified common surface protein regulators. a**, Venn diagram of all genes enriching (lfc > 2; p-val < 0.01) in at least one of four screens. **b**, GO-term analysis of genes from (b) enriching in >=2 screens. BP = biological process, CC = cellular component. **c**, String-DB network based on interaction confidence (medium-confidence setting) of all genes shown in (a) enriching in >= 2 screens. Selected complexes or pathways were manually annotated. **d**, Results from FACS-based CRISPR screens targeting indicated markers, split by sampling timepoint. Genes highlighted in red are strongly depleting in this screen (LFC vs t0 < -4).

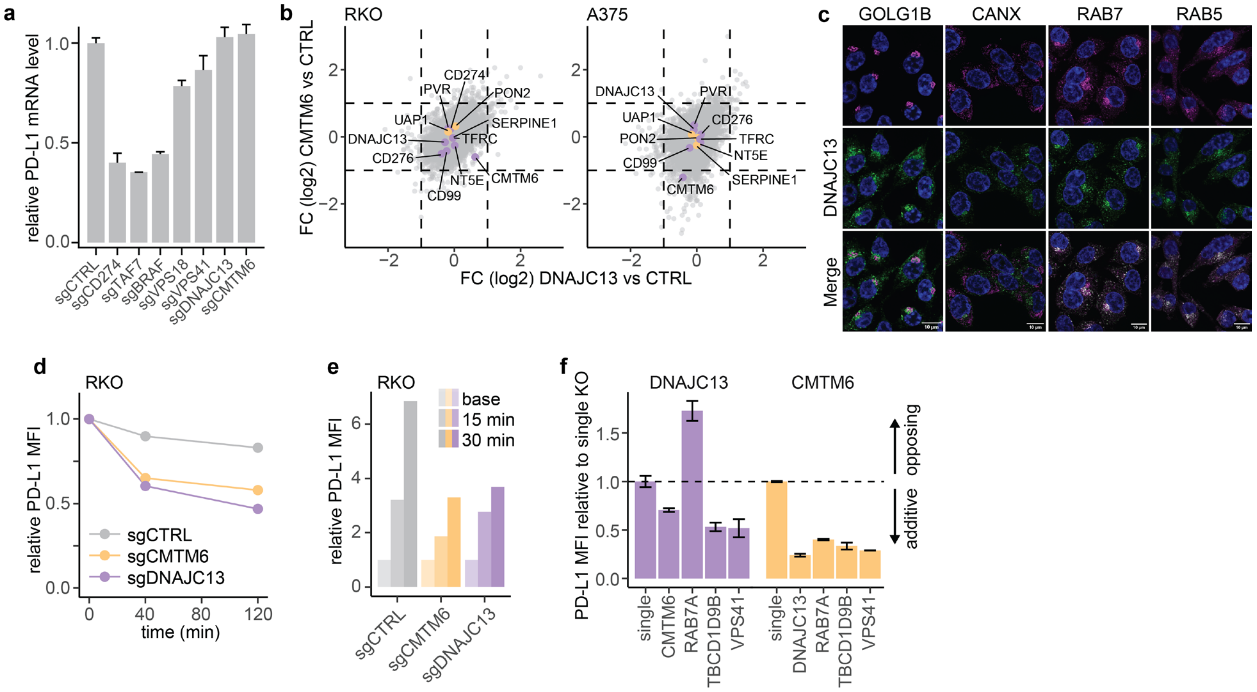


**Extended data Fig. 6 | DNAJC13 is a post-translational mediator of PD-L1 expression. a**, qPCR of CD274 mRNA 4 days after inducible KO of selected genes. Mean of 2 independent replicates with different guide RNAs is shown with error bars indicating the positive standard deviation. **b**, mRNA from RKO cells (left) or A375 cells (right) harboring perturbations of DNAJC13, CMTM6 or control was isolated 10 days after CRISPR editing and sequenced (Lexogen Quantseq). **c**, Localization of endogenous DNAJC13 was assessed using markers for the Golgi apparatus (GOLG1B), Endoplasmatic reticulum (CANX), and endosomal compartment (RAB7 for late endosomes and RAB5 for early endosomes). Scale bar 10µM. **d**, RKO cells were stained with unconjugated IgG-anti-PD-L1 antibody, thoroughly washed and then placed at 37 degrees. PD-L1 MFI was assessed by anti-IgG staining at indicated timepoints. **e**, RKO cells were stained with unconjugated IgG-anti-PD-L1 antibody, thoroughly washed and then placed at 37 degrees for 60 minutes to allow internalization. Cells were then washed with a low pH stripping buffer to remove surface bound antibody. Cells were again placed on 37 degrees to allow re-expression of internalized (and stripping-protected) antibody-bound PD-L1. At indicated timepoint, re-expression was assessed by anti-IgG staining and flow-cytometry. **f**, Combinatorial perturbation of DNAJC13 or CMTM6 and selected trafficking factors. Effects were normalized to single perturbations (sgDNAJC13 or sgCMTM6), n = 3.

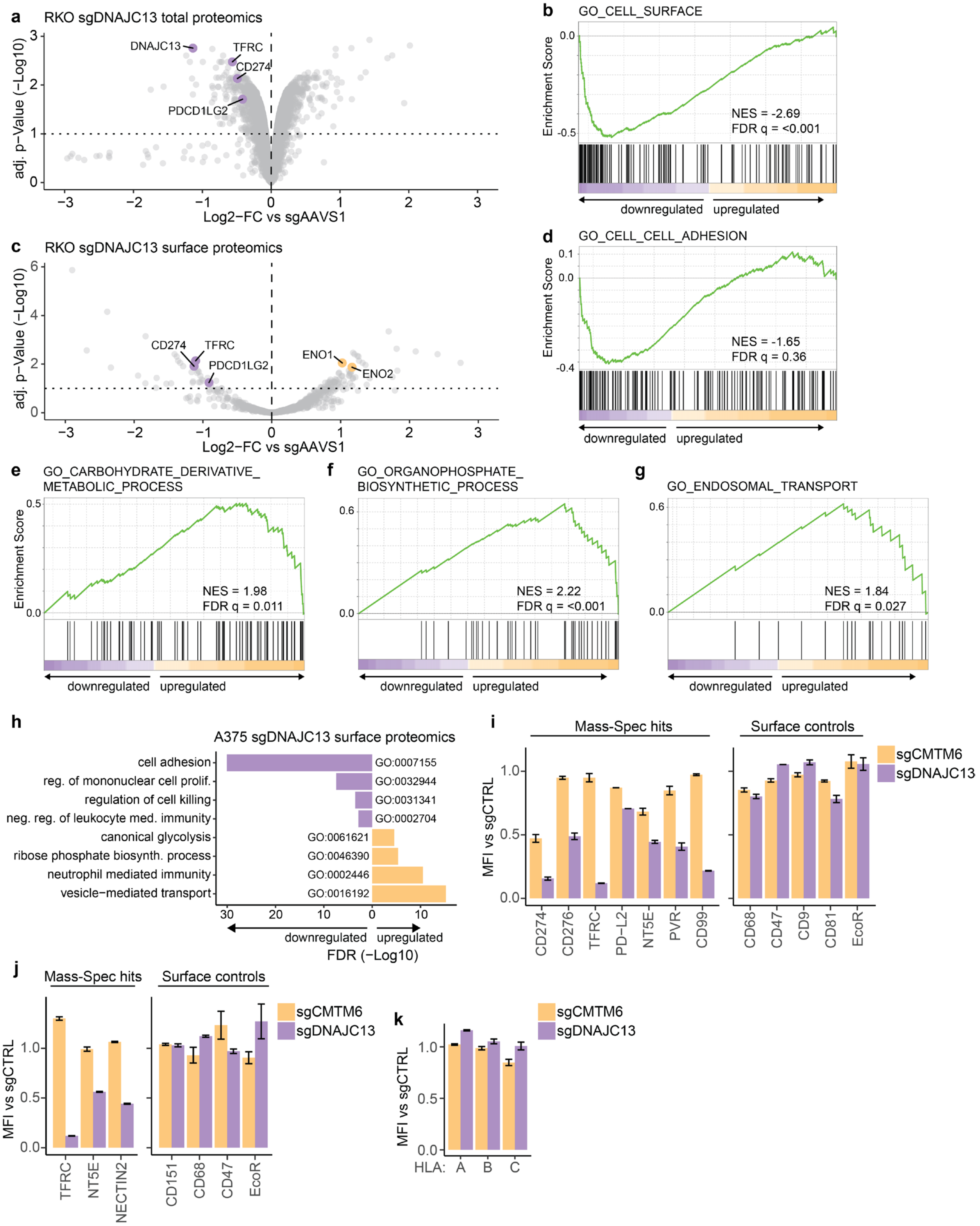


**Extended data Fig. 7 | Proteomics of sgDNAJC13 KO cells. a**, Total proteomics in RKO cells expressing sgDNAJC13 comparing to sgAAVS1. Cells were cultured for 10 days after inducible editing. **b**, Rank-based Gene Set Enrichment Analysis (GSEA) of significantly (adj. p < 0.1) altered proteins in sgDNAJC13+ RKO cells. **c**, Surface proteomics in RKO cells expressing sgDNAJC13 comparing to sgAAVS1. Cells were cultured for 10 days after inducible editing. **d**, Rank-based GSEA of altered surface proteins in sgDNAJC13+ RKO cells. **e – f**, Rank-based GSEA of sgDNAJC13+ A375. For all GSEA shown, Log2-FC vs. sgAAVS1 were used to compute enriched gene sets (GO biological process). NES = Normalized enrichment score. **h**, GO-term (biological process) enrichment analysis of significantly de-regulated surface proteins (adj. p-Value < 0.1) in sgDNAJC13 expressing A375 cells. Analysis was performed using PANTHER with Fisher's exact test and calculation of false discovery rate. **i-j**, Flow cytometry validation of Mass-Spec hits and other expressed surface proteins in RKO (i) and A375 (j) cells. MFI of respective markers was compared to Control KO cells. EcoR = murine Slc7a1, used in Cas9 clone engineering. **k**, HLA class I expression in A375 CMTM6 or DNAJC13 KO cells. MFI of PD-L1 was compared to Control KO cells (see also new extended Figure 7k). For i-j, n = 2 or 3 independent replicates.

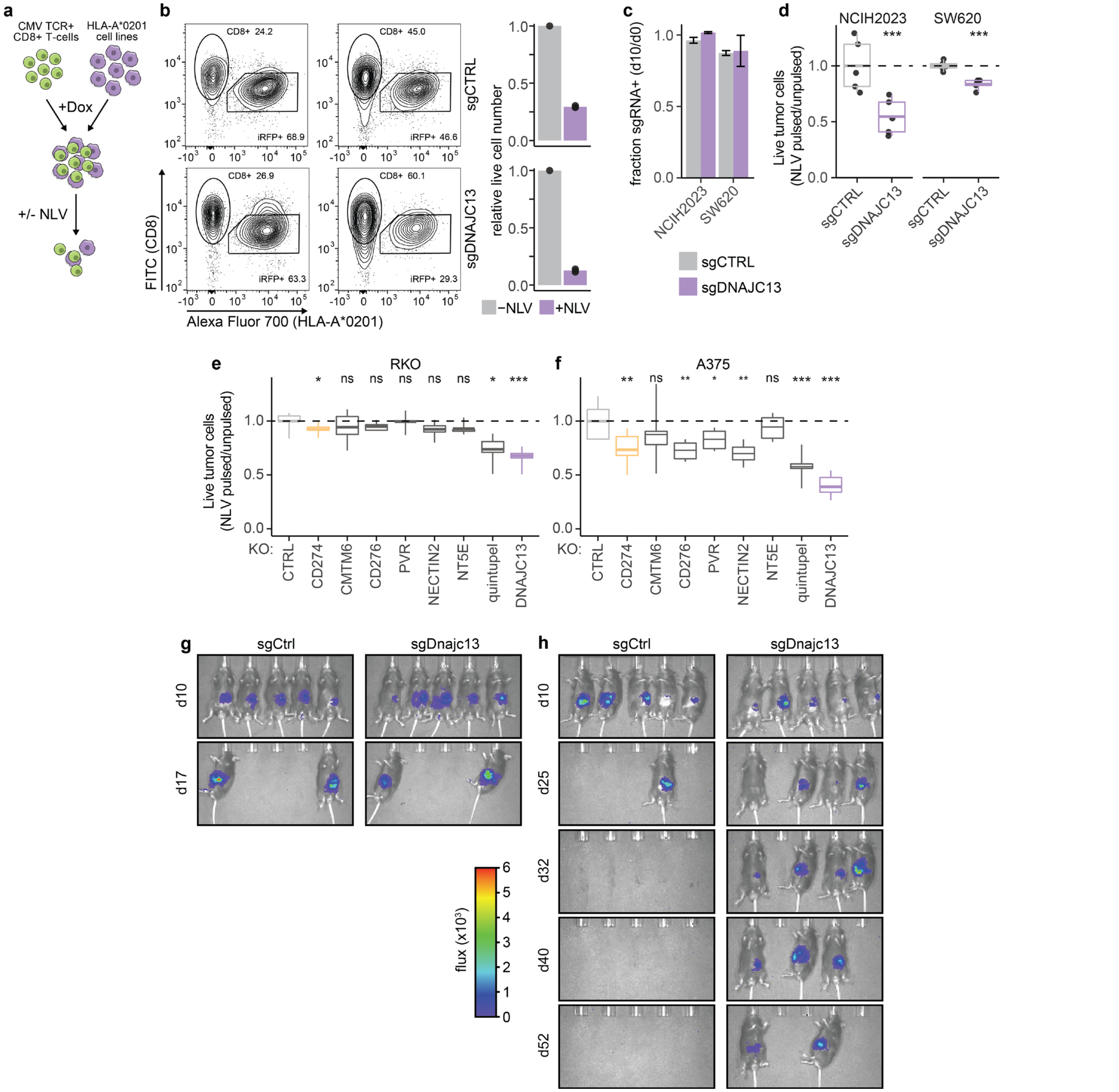


**Extended data Fig. 8 | Suppression of DNAJC13 enhances CD8+ T cell-mediated tumor cell killing. a**, Schematic of co-culture assays to measure anti-tumor effects of human CD8⁺ T cells in diverse human cell lines. **b**, Representative FACS plots and quantification of RKO – CD8⁺ T-cell co-culture assays, gated on live cells. **c**, Fitness effect upon KO of DNAJC13 in cancer cell lines from different tissue contexts. Cells transduced with indicated sgRNAs were cultured for 10 days following Cas9 induction, and the fraction of sgRNA+ cells was quantified relative to the initial population prior to Cas9 induction. **d-f**, Comparison of live tumor cell counts in NLV-pulsed samples and unpulsed conditions in NCIH2023 (d, left) or SW620 (d, right), RKO (e), A375 (f) cells. NCIH2023, SW620, A375 were pre-treated with 10ng/ml IFN-y 48h prior to addition of T-cells. Effect sizes were normalized to the respective sgCTRL sample. Boxplots show individual values from replicates (n = 5-8) with whiskers indicating Min. to Max. ‘quintupel’ cells harbor deletions of CD274, CD276, PVR, NECTIN2 and NT5E. p-values were calculated using unpaired t-tests with Welch correction, with * = $p < 0.05$, ** = $p < 0.005$, *** < $p = 0.0005$. **g-h**, Luciferin bioluminescence imaging of Rag2-/- (g) or WT (h) B6 mice upon orthotopic transplantation of sgCtrl or sgDnajc13 transduced EPP2 pancreas adenocarcinoma cells. Representative images of experiment shown in Figure 6c.

### Materials and Methods

#### Table 1: FACS antibodies

| Antibody | clone | fluorophore | Dilution used | Source |
|---|---|---|---|---|
| PD-L1 | 29E.2A3 | PE-Cy7 | 1:400 | Biolegend, 329718 |
| CD47 | B6H12 | APC | 1:400 | eBioscience, 14-0479-82 |
| CD276 | DCN.70 | PE | 1:400 | Biolegend, 331606 |
| CD151 | 50-6 | APC | 1:400 | Biolegend, 350406 |
| HLA-E | 3D12 | PE-Cy7 | 1:50 | Biolegend, 342608 |

#### Table 2: Plasmids

| **Plasmid name** | **Purpose** |
|---|---|
| pWXLd-EF1αs-rtTA3-IRES-EcoR-PGK-PuroR | Inducible Cas9 |
| pLenti v2-TRE3G-Cas9-P2A-GFP-PGK-BlastR | Inducible Cas9 |
| pRRL-SFFV-rtTA3-IRES-EcoR-PGK-HygroR | Inducible Cas9 |
| pLenti v2-TRE3G-Cas9-P2A-BFP | Inducible Cas9 |
| pRRL-PBS-U6-sgRNA-EF1αs-Thy1-P2A-NeoR | sgRNA library |
| pRRL-U6-sgRNA-EF1αs-mCherry-P2A-NeoR | Single sgRNA |
| pLenti v2-U6-sgRNA-PGK-GFP-P2A-NeoR | Single sgRNA |
| SFFV-TCRa-P2A-TCRb | CMV specific TCR |
| SFFV-HLA-A*0201-IRES-iRFP720 | T-cell co-cultures |

**sgRNA cloning**

sgRNA oligos were ordered from Sigma-Aldrich and annealed and phosphorylated using T4 PNK (NEB). sgRNAs were cloned in Lenti-hU6-sgRNA-improved tracer-EF1a-mCherry-P2A-NeoR or Lenti-hU6-sgRNA-improved tracer-EF1a-GFP-P2A-NeoR vector by overnight digestion of the vector backbone with BsmBI, followed by ligation with annealed sgRNA oligos.

**Lentiviral Transfection and transduction of cells**

Lenti-X 293T cells (Clontech, 632180) were transfected upon being 70-80% confluent with Lentiviral plasmid, packaging plasmid pCMVR8.74 (addgene, 22036) and envelope plasmid pCMV-VSVG (addgene, 8454) at a ratio of 4:2:1 in serum-free DMEM. PEI was added and incubated for 20 minutes at RT before adding dropwise to cells. Medium was changed to target cell medium after 12-18 hours. Supernatants were harvested several times between 48 and 72 hours after transfection and filtered through 0.45um filter. Target cells were transduced with the viral supernatant with optional addition of 4ug/ml polybrene (Sigma-Aldrich, H9268). Leftover virus was concentrated using PEG, aliquoted and frozen.

**Cell culture and iCas9-clone generation**

Human cancer cell lines were acquired from ATCC. A375 cells were a gift from Anna Obenauf (IMP Vienna). Cells were routinely cultured in either DMEM or RPMI-1640 (Gibco) supplemented with 10% FBS (Sigma-Aldrich), 20 mM L-glutamine, 10 mM sodium pyruvate, 100 U/ml penicillin and 100 µg/ml streptomycin. Cell lines were transduced with either rtTA construct and Tet-inducible Cas9 vector and single cell sorted upon doxycycline induction. Growing clones were tested for functional Cas9 after transduction with sgRNAs targeting an essential gene (PLK1 in Human cells and Rpa3 in mouse cells). Clones were selected based upon depletion of the transduced population in a FACS based competition assay upon addition or no addition of doxycycline.

**Pooled sgRNA screening**

For the genome-wide screening, more than one billion cells, divided in two replicates were transduced with the sgRNA library lentivirus at an infection efficiency of 15% resulting in 500x representation of the library for each of the replicate.

Cells were selected on 1 mg/ml G418 before harvesting for a day 0 timepoint and induction of Cas9 with 500 ng/ml doxycycline (T0) for 2 days. Leftover G418 selected cells were frozen for future use, from which about 250 million cells per replicate were thawed for a second round of screens (CD47, CD276, HLA-E and CD151). Transduced cells were counted and passaged every two days until day 18 when cells were again harvested for DNA extraction. In all cases, on day 4 and day 10 after Cas9 induction, cells were stained with respective antibodies and sorted on BD FACSAria III cell sorters for the POI-low population. 200 million cells per sample were stained with respective antibodies (Table 1) in 10 ml staining mix (PBS, 5% FBS, 2mM EDTA). For all samples, the low 1% population of aPOI was sorted, except for PD-L1 day 10 for which the lower 0.1% population was sorted and for HLA-E for which lower 2% was sorted at both time points.

NGS libraries were prepared as previously described[15] with minor modifications. Briefly, sgRNA cassettes were amplified using a nested PCR to first isolate the sgRNA sequence from genomic DNA and then to add sample barcodes and sequencing adapters compatible with Illumina's HiSeqV4. The first PCR reaction was performed using 28 cycles, 52ºC annealing temperature and using 1000 ng template per reaction (for POI neg samples as low as 100 ng per reaction). Using gel purified (or column purified for POI negative samples) product from the first PCR reaction, the secondary PCR was performed using 6 cycles, 57ºC annealing and 10 ng of the template per reaction. Libraries were multiplexed using indexes compatible with the Illumina TruSeq HT kit with the primers below where N denotes a six basepair index. All libraries were sequenced on a HiSeq2500 platform (Illumina).

To quantify raw sequencing reads we used the *crispr-process-nf* Nextflow workflow available at https://github.com/ZuberLab/crispr-process-nf. Briefly, all guides in the sgRNA library were padded with Cs to equal length before creating an index for bowtie2[15,55]. Random 6mer nucleotides were trimmed using the fastx_trimmer from the fastx-toolkit v0.0.14 (http://hannonlab.cshl.edu/fastx_toolkit/) before demultiplexing via 4mer sample barcodes with fastx_barcode_splitter (--mismatches 1 --bol). Next, barcodes and 20mer spacer were trimmed using fastx_trimmer and reads were aligned with bowtie2 v2.3.0 (-L 18 --score-min 'C,0,-1' -N 0 --seed 42) and quantified with featureCounts v1.6.1[56]. To calculate enrichment or depletion of sgRNAs we implemented the *crispr-mageck-nf* Nextflow workflow available at https://github.com/ZuberLab/crispr-mageck-nf. First, count tables were filtered to exclude sgRNAs with less than 50 counts in control samples prior to further downstream analyses. Read counts were median normalized and average log2 fold-changes, p-values and FDRs were calculated using MAGeCK 0.5.9[57]. To calculate enrichment of sgRNAs in sorted populations, sgRNA counts within the sorted populations were compared to the T0 populations.

**Cell Line Authentication**

Short tandem repeat (STR) profiles were determined for each line using the Promega PowerPlex 16 System (performed in-house at IMP). STR profiles were compared with external STR profiles of cell lines (ATCC or DSMZ databases). All stocks were tested for mycoplasma contamination before and after cells were cryopreserved using real-time PCR based method, using primers specific to detect 220 different species.

**Recycling assays**

A375 or RKO cells were stained with unconjugated IgG-anti-PD-L1 antibody, thoroughly washed and then placed at 37 degrees to allow internalization and recycling/degradation. PD-L1 MFI was assessed over time by anti-IgG staining at indicated timepoints. For re-expression assays, cells were stained with unconjugated IgG-anti-PD-L1 antibody, thoroughly washed and then placed at 37 degrees for 60 minutes to allow internalization. Cells were then washed twice with a low pH stripping buffer (0.5 M NaCl, 0.5% acetic acid, pH 2.5–2.8) at 4 degrees to remove surface bound antibody. Cells were again placed on 37 degrees to allow re-expression of internalized (and therefore stripping-protected) antibody-bound PD-L1. At indicated timepoint, re-expression was assessed by anti-IgG staining and flow-cytometry.

**Mass spectrometry based total and surface proteomics**

*Total proteomics*

RKO and A375 iCas9 clones were lentivirally transduced with sgRNAs and cultured for 10 days after Cas9 induction, washed twice with PBS and pelleted. Each cell pellet was lysed in 520 µl of 10M Urea and 50 mM HCl and incubated 10 min at room temperature before addition of 62.5 µl 1M Tris (pH 8) The amount of protein was determined in a Bradford assay. Each sample was incubated with 250 units Benzonase (250 U/µl, purity grade I, Merck) and 5 µl 1M

Dithiothreitol (DTT) at 37°C for 1h. The alkylation was performed by adding 10 µl of 1M Iodoacetamide and incubating for 30 min in the dark. The reaction was quenched by addition of 2.5 µl 1M DTT. Samples were diluted to 6M urea by addition of 100mM Tris pH 8 and the proteins were digested with Lys-C at an enzyme to protein ration of 1:50 for 3h at 37ºC. Subsequently, the samples were further diluted to 2M urea with 100 mM TRIS pH 8 and a tryptic digest was done overnight at 37ºC with an enzyme to protein ratio of 1:50. The samples were acidified by addition of 10% Trifluoroacetic acid to reach pH 2. Peptides were desalted using C18 cartridges (Sep-Pak Vac 1cc (50mg), Waters). Peptides were eluted with 2 x 200 µl 80% Acetonitrile (ACN) and 0.1% Formic Acid (FA), followed by freeze-drying.

Labelling with TMT: The peptides were dissolved in 100 µl 100mM Triethylammoniumbicarbonate (TEAB). 100 µg (in 50 µl) of each sample were labelled with one separate channel of the TMT 10 plex reagent (Thermo Fisher) according to the manufacturer's description. The labelling efficiency was determined by LC-MS/MS on a small aliquot of each sample. Samples were mixed in equimolar amounts and equimolarity was again evaluated by LC-MS/MS. The sample was acidified to a pH below 2 with 10% TFA and was desalted as described above.

SCX-separation: The dried sample was dissolved in 220 µl of SCX Buffer A (5 mM NaH2PO4, pH 2.7, 15% ACN) and 200µg of peptide were loaded on the column. SCX was performed using a custom-made TSKgel SP-2PW SCX column (5 µm particles, 12.5 nm pore size , 1 mm i.d. x 250 mm, TOSOH) on an Ultimate system (Thermo Fisher Scientific) at a flow rate of 35 µl/min. For the separation, a ternary gradient was used. Starting with 100% buffer A for 10 min, followed by a linear increase to 10% buffer B (5 mM NaH2PO4, pH 2.7, 1M NaCl, 15% ACN) and 50% buffer C (5 mM Na2HPO4, pH 6, 15% ACN) in 80 min, to 25% buffer B and 50% buffer C in 10 min, 50% buffer B and 50% buffer C in 10 min and an isocratic elution for further 15 min. The flow-through was collected as single fraction, along the gradient fractions were collected every minute. 120 fractions were collected, ACN was removed by vacuum centrifugation and the samples were acidified with 0.1% TFA and analyzed by LC-MS/MS.

LC-MS/MS: 5% to 30% of each SCX -fraction were analysed by LC-MS/MS. The nano HPLC system used was a Thermo Fisher RSLC nano system (Thermo Fisher Scientific, Amsterdam, Netherlands) coupled to a Q Exactive HF mass spectrometer (Thermo Fisher Scientific, Bremen, Germany), equipped with a Proxeon nanospray source (Thermo Fisher Scientific, Odense, Denmark). Peptides were loaded onto a trap column (Thermo Fisher Scientific, Amsterdam, Netherlands, PepMap C18, 5 mm × 300 µm ID, 5 µm particles, 100 Å pore size) at a flow rate of 25 µL min-1 using 0.1% TFA as mobile phase. After 10 min, the trap column was switched in line with the analytical column (Thermo Fisher Scientific, Amsterdam, Netherlands, PepMap C18, 500 mm × 75 µm ID, 2 µm, 100 Å). The gradient starts with the mobile phases: 98% A (water/formic acid, 99.9/0.1, v/v) and 2% B (water/acetonitrile/formic acid, 19.92/80/0.08, v/v/v), increases to 35%B over the next 60min, followed by a gradient in 5 min to 90%B, stays there for five min and decreases in 5min back to the gradient 98%A and 2%B for equilibration at 30°C. The Q Exactive HF mass spectrometer was operated in data-dependent mode, using a full scan (m/z range 350-1650, nominal resolution of 120,000, target value 3E6) followed by MS/MS scans of the 10 most abundant ions. MS/MS spectra were acquired using normalized collision energy of 35%, isolation width of 1.2 m/z, resolution of 60.000 and the target value was set to 1E5 and first fixed mass is set to 115 m/z. Precursor ions selected for fragmentation (exclude charge state 1, 7, 8, >8) were put on a dynamic exclusion list for 30 s. Additionally, the minimum AGC target was set to 1E5 and intensity threshold was calculated to be 1E4. The peptide match feature was set to preferred and the exclude isotopes feature was enabled.

Data processing: For peptide identification, the RAW-files were loaded into Proteome Discoverer (version 1.4.0.288, Thermo Scientific). All hereby created MS/MS spectra were searched using MSAmanda v1.4.14.8240[58]. The RAW-files were searched against the human SwissProt database (release 2017_09) with common contaminants appended. The following search parameters were used: Carbamidomethylation of cysteine and 10-plex tandem mass tag® (TMT) on lysine and N-termini were set as fixed modification and oxidation of methionine was set as variable modification. Monoisotopic masses were searched within unrestricted protein masses for tryptic enzymatic specificity. The peptide mass tolerance was set to ±5 ppm and the fragment mass tolerance to ±0.03 Da. The maximal number of missed cleavages was set to 2. The result was filtered to 0.5% FDR on peptide spectrum match level using Percolator algorithm integrated in Thermo Proteome Discoverer. Peptides were quantified based on Reporter Ion intensities extracted by the "Reporter Ions Quantifier"-node implemented in Proteome Discoverer. Statistical significance of differentially expressed proteins were determined using limma[59].

*Surface proteomics*

RKO and A375 iCas9 clones were transduced with sgAAVS1, sgCMTM6 or sgDNAJC13 and cultured for 10 days after Cas9 induction. A375 were pre-treated with 10ng/µl IFNy for 48 hours. Surface-protein enrichment followed by label-free mass spectrometry was performed as previously described[60] with the modification that trypsin-resistant Neutravidin-beads were used[61].
Raw files were processed using MaxQuant (version 1.5.1.2). Spectra were searched against the UniProt database consisting of reviewed human proteins (June 2020). The Andromeda search engine was used with the following search criteria: enzyme was set to trypsin/P with up to 2 missed cleavages. Carbamidomethylation (C) was selected as a fixed modification; oxidation (M), acetylation (protein N-term) were set as variable modification. Match between runs was enabled with match time window set to 1 min and alignment time window set to 20 min. The search type for protein quantification was set to standard. Quantification intensities were calculated by the default fast MaxLFQ algorithm with minimal ratio count set to 2. Require MS/MS for LFQ comparisons was disabled. Peptide and protein hits were filtered at a false discovery rate of 1%, with a minimal peptide length of 7 amino acids. Second peptide search for the identification of chimeric MS2 spectra was enabled. Not mentioned MaxQuant settings were left as default.
Mass spectrometry raw data is available on ProteomeXchange via the PRIDE database, ID PXD021989.

**Human T-cell co-cultures**

Human CD8+ T cells were obtained from Stem Cell (#70027). CD8 expression was confirmed by flow-cytometry. pMP71-CMV.NLV-TCR1 was a gift from Ton Schumacher and the TCRalpha-P2A-TCRbeta cassette was PCR amplified and cloned into pRRL-SFFV-BamHI-SalI-WPRE after BamHI/SalI digest using Gibson assembly. CD8+ T cells were transduced with the CMV.NLV-TCR1 construct by lentiviral delivery and the positive population enriched by FACS sorting. T cells were cultures in RPMI-1640 (Gibco), supplemented with 10% FBS (Sigma-Aldrich), 20 mM L-glutamine, 10 mM sodium pyruvate, 1x MEM NEAA solution (Gibco), 100 U/ml penicillin and 100 µg/ml streptomycin, 50 uM b-Mercaptoethanol, 20mM HEPES.
A codon optimised cDNA sequence of HLA-A*0201 (sequence source: pCIpA102-G-HLA-A2_GFP, addgene) was PCR amplified from a GeneBlock (Integrated DNA Technologies).
iCas9-RKO cells were first transduced with SFFV-HLA-A*0201-IRES-iRFP720 and sorted for iRFP720 expressing cells by FACS. Other iCas9 cell lines used harbor intrinsic HLA-A*0201 alleles. HLA-A*0201 expression was confirmed by flow cytometry. 50,000 iCas9-RKO, -A375, -MDAMB231, 639V or NCIH2023 cells, or 100,000 SW620 cells, were seeded in 96-well plates and selected wells pulsed with 10 ng/ml NLV peptide (IBA Lifesciences, #6-7001-901) at 37°C for at least 2 hours. Supernatant was removed, cells were washed and either 25,000 or 50,000 CD8+ T cells were added in T cell growth medium. After 24 or 48 hours, the entire well was harvested (including all wash supernatants) and cells stained for viability and subsequently analyzed by flow cytometry (iQue Screener Plus, Sartorius).

**Mouse experiments**

Mice were bred and maintained in the IMP/IMBA facility in standard pathogen-free and controlled conditions: temperature at 22 ± 1 °C, humidity at 55 ± 5 %, and a 14-hour light/10-hour dark cycle, with unrestricted access to food and water. All animal experiments were conducted in compliance with protocols approved by the Austrian Ministry (GZ: 2260492-2022-22). The generation of the EPP2 pancreas adenocarcinoma cell line and injection methods were previously reported[49]. Briefly summarized, mice were anesthetized with isoflurane and placed on a heated surface. A small incision was made in the upper left abdomen after disinfection, the spleen located, and the pancreas gently externalized. A solution of cells in PBS (5x105 cells in 10 µl) was injected into the pancreas using a Hamilton syringe. After injection, internal organs were carefully repositioned, the muscle incision site was sutured with resorbable 6-0 Vicryl, and the skin closed with sterile wound clips. Mice were administered 5 mg/kg carprofen intraperitoneally as preemptive analgesia and again every 12 h for 48 h post-operation. Animal health was monitored daily.

**H3K27ac ChIP-seq Re-analysis and Super-enhancer Analysis**

All ChIP-seq data were aligned to the human genome assembly GRCh37/hg19 using Bowtie2 version 2.3.5.1. The resulting SAM file was converted to a BAM file using SAMtools version 1.9 and used for peak finding. To identify super-enhancer regions in RKO cells similar to Whyte et al.[62] and Nakamura et al.[63] by exceptionally broad regions of enriched active chromatin, H3K27ac ChIP-seq data (SRX2631830) from Nakamura et al.[63] and the corresponding input sample (SRX2631831) were analyzed using the MACS2 algorithm version 2.1.2 with the following settings: Broad region calling on; p-value cutoff for broad regions = 0.1; p-value cutoff for narrow/strong regions = 0.01; band width 300; without building the shift model and 200 bp fragment extension. The resulting 97889 peak regions were overlapped using Multovl version 1.4[64] with 411 ENCODE blacklist regions for the human genome assembly GRCh37/hg19 (https://www.encodeproject.org/files/ENCFF001TDO), leaving 97759 peaks, of which 21402 peaks with a FDR <= 0.01 were used for plotting. H3K27ac ChIP-seq data of MIAPACA2 cells (SRX4645325), generated by Somerville et al.[65], serve as a reference for the *CD274* locus in Extended data Fig. 1). GRCh37/hg19 mapped and RPM-normalized bigwig files of the indicated SRX tracks from www.https://chip-atlas.org/ were used for ChIP-seq data visualization in IGV version 2.9.4.
